# Unique Transcription Factor Functions Regulate Epigenetic and Transcriptional Dynamics During Cardiac Reprogramming

**DOI:** 10.1101/642900

**Authors:** Nicole R. Stone, Casey A. Gifford, Reuben Thomas, Karishma J. B. Pratt, Kaitlen Samse-Knapp, Tamer M. A. Mohamed, Ethan M. Radzinsky, Amelia Schricker, Pengzhi Yu, Kathryn N. Ivey, Katherine S. Pollard, Deepak Srivastava

**Author notes:** Correspondence to: Deepak Srivastava Gladstone Institutes, 1650 Owens St., San Francisco, CA 94158, Katherine S. Pollard Gladstone Institutes 1650 Owens St., San Francisco, CA 94158. co-first authors (Nicole R. Stone, Casey A. Gifford).

## Abstract

Direct lineage conversion, whereby a somatic cell assumes a new cellular identity, can be driven by ectopic expression of combinations of lineage-enriched transcription factors. To determine the molecular mechanisms by which expression of Gata4, Mef2c, and Tbx5 (GMT) induces direct reprogramming from a cardiac fibroblast toward an induced cardiomyocyte, we performed a comprehensive transcriptomic and epigenomic interrogation of the reprogramming process. Single cell RNA sequencing indicated that a reprogramming trajectory was acquired within 48 hours of GMT introduction, did not require cell division, and was limited mainly by successful expression of GMT. Evaluation of chromatin accessibility by ATAC-seq supported the expression dynamics and revealed widespread chromatin remodeling at early stages of the reprogramming process. Chromatin immunoprecipitation followed by sequencing of each factor alone or in combinations revealed that GMT bind DNA individually and in combination, and that ectopic expression of either Mef2c or Tbx5 is sufficient in some contexts to increase accessibility. We also find evidence for cooperative facilitation and refinement of each factor’s binding in a combinatorial setting. A random-forest classifier that integrated the observed gene expression dynamics with regions of dynamic chromatin accessibility suggested Tbx5 binding is a primary driver of gene expression changes and revealed additional transcription factor motifs co-segregating with reprogramming factor motifs, suggesting new factors that may be involved in the reprogramming process. These results begin to explain the mechanisms by which transcription factors normally expressed in multiple germ layers can function combinatorially to direct lineage conversion.

## INTRODUCTION

Somatic cellular identity is established by complex gene regulatory networks during embryonic development. Cellular reprogramming, whereby ectopic expression of transcription factors promotes acquisition of a new cell state, challenges the notion that somatic identity is permanent (Guo and Morris, 2017). Knowledge regarding molecular mechanisms that dictate embryonic development has been exploited to devise combinations of transcription factors that can facilitate direct reprogramming of fibroblasts toward myriad somatic cell types, including macrophages, neurons, and cardiomyocytes, without progression through an intermediate pluripotent state (Feng et al., 2008; Ieda et al., 2010; Rackham et al., 2016; Wernig et al., 2008). With a few exceptions (Davis et al., 1987; Xie et al., 2004), cellular reprogramming typically requires a combination of three to five lineage-enriched, but not lineage-specific, factors, although the precise mechanisms by which various combinations lead to cellular specificity is only beginning to be understood (Treutlein et al., 2016; Wapinski et al., 2017). Beyond offering a promising alternative to induced pluripotency to create resources for cell therapy, cellular reprogramming offers a unique system with which to study transcription factor function (Wapinski et al., 2013). The heterogeneity, functional redundancy, and transient nature of developing tissues has made mechanistic studies aimed at dissecting individual protein functions and the interdependency of transcription factor binding difficult to ascertain.

Direct reprogramming of cardiac fibroblasts to cardiomyocytes has been achieved by ectopic expression of cardiac-enriched factors (Fu et al., 2013; Ieda et al., 2010; Nam et al., 2013; Qian et al., 2012; Song et al., 2012). Ectopic expression of Gata4, Mef2c, and Tbx5 (GMT) is sufficient to alter the fibroblast epigenome and promote expression of genes associated with cardiomyocytes while simultaneously repressing the fibroblast gene program (Ieda et al., 2010; Liu et al., 2017; Zhou et al., 2016). Perturbation of epigenetic remodelers and ectopic expression of additional transcription factors such as Hand2 and MYOCD were found to influence reprogramming as well, with addition of Hand2 resulting in a greater portion of pacemaker-like cells (Addis et al., 2013; Christoforou et al., 2013; Protze et al., 2012; Zhou et al., 2016). Concomitant inhibition of TGF**β** and Wnt signaling resulted in improved reprogramming both *in vitro* and *in vivo* (Ifkovits et al., 2014; Mohamed et al., 2017). Despite the relatively inefficient nature of this process, regardless of approach, *in vivo* studies have shown that reprogramming offers a therapeutic benefit (Jayawardena et al., 2015; Qian et al., 2012; Song et al., 2012).

Gata4, Mef2c, and Tbx5 each have essential functions in a wide range of tissues during embryonic development. Deletion of any of these factors individually leads to murine embryonic lethality by E10.5 and gross malformation of various organ systems, including the developing cardiovascular system (Bruneau et al., 2001; Lin et al., 1997; Molkentin et al., 1997). Gata4 is a zinc finger transcription factor capable of interacting with compact chromatin, but exhibits limited ability to interact with regions containing DNA methylation (Cirillo et al., 2002; Oda et al., 2013). While it interacts with Nkx2-5 and Tbx5 to promote cardiovascular development, it also cooperates with Foxa2 during endoderm development to promote expression of a transcriptional network required for foregut development (Holtzinger and Evans, 2005). Similarly, beyond cardiogenesis Tbx5 is also required for limb development (Agarwal et al., 2003; Ahn et al., 2002), while Mef2c is also essential for neural development (Akhtar et al., 2012; Flavell et al., 2006; Leifer et al., 1993; Li et al., 2008a, 2008b; Shalizi et al., 2006). Thus, transcription factors must interact in a combinatorial fashion to induce genome-wide epigenetic changes, but the mechanism by which these factors achieve tissue specific DNA binding patterns and transcriptional regulation remains unknown.

Although Gata4 in other contexts can interact with heterochromatic genomic regions that are relatively inaccessible, a necessary event during reprogramming, little is known regarding the ability of Tbx5 and Mef2c to bind to closed areas of chromatin. One screen identified TBX5 as a factor capable of inducing DNA demethylation when expressed ectopically (Suzuki et al., 2017). While the function of Mef2c in this regard remains unclear, a study on the closely related factor Mef2d in photoreceptor cells found that it requires additional co-factors to access regions that do not encode strong consensus Mef-response motifs (Andzelm et al., 2015). Which of these factors, if any, function to open closed chromatin in the context of direct reprogramming, and whether they require combinatorial interaction to do so, remains unknown.

Here, we investigated the genome-wide consequences of Gata4, Mef2c and Tbx5 expression, alone and in combination, in cardiac fibroblasts as the cells underwent reprogramming towards a cardiomyocyte-like state. By combining single cell RNA sequencing, ChIP-seq, and ATAC-seq analyses, we found that epigenomic and transcriptional changes occurred rapidly within the first 24-48 hours of reprogramming. Cells that adopted a trajectory toward the cardiac fate could largely be predicted by virtue of early gene expression changes and reprogramming factor expression. Applying a machine learning approach identified new candidate factors involved in reprogramming. Although GMT are each capable of acting independently to promote chromatin remodeling when expressed individually, we found that changes were primarily associated with Mef2c and Tbx5 binding only. Cooperative activity between Gata4, Tbx5, and Mef2c was evident as combinatorial expression resulted in refinement and facilitation of DNA binding compared to single factor expression, and combinatorial binding correlated with opening of chromatin particularly at cardiac loci.

## RESULTS

### Multiple Transcription Signatures Identified During Cardiac Reprogramming

To determine the discrete temporal transcriptional response to reprogramming with GMT in the setting of TGF**β** and Wnt inhibition, we performed single cell RNA sequencing during cardiac reprogramming of Thy1 positive (Thy1+) cells, largely representing fibroblasts, isolated from neonatal mouse hearts that encode an αMHC-GFP reporter that is activated during reprogramming (Ieda et al., 2010) (**Figure 1A**). We collected and analyzed 29,718 cells representing five time points after transduction with retroviruses encoding Gata4, Mef2c, and Tbx5 (days 1, 2, 3, and 7), as well as the starting population. We additionally collected cells sorted at day 14 using the aforementioned αMHC-GFP reporter.

**Figure 1.**
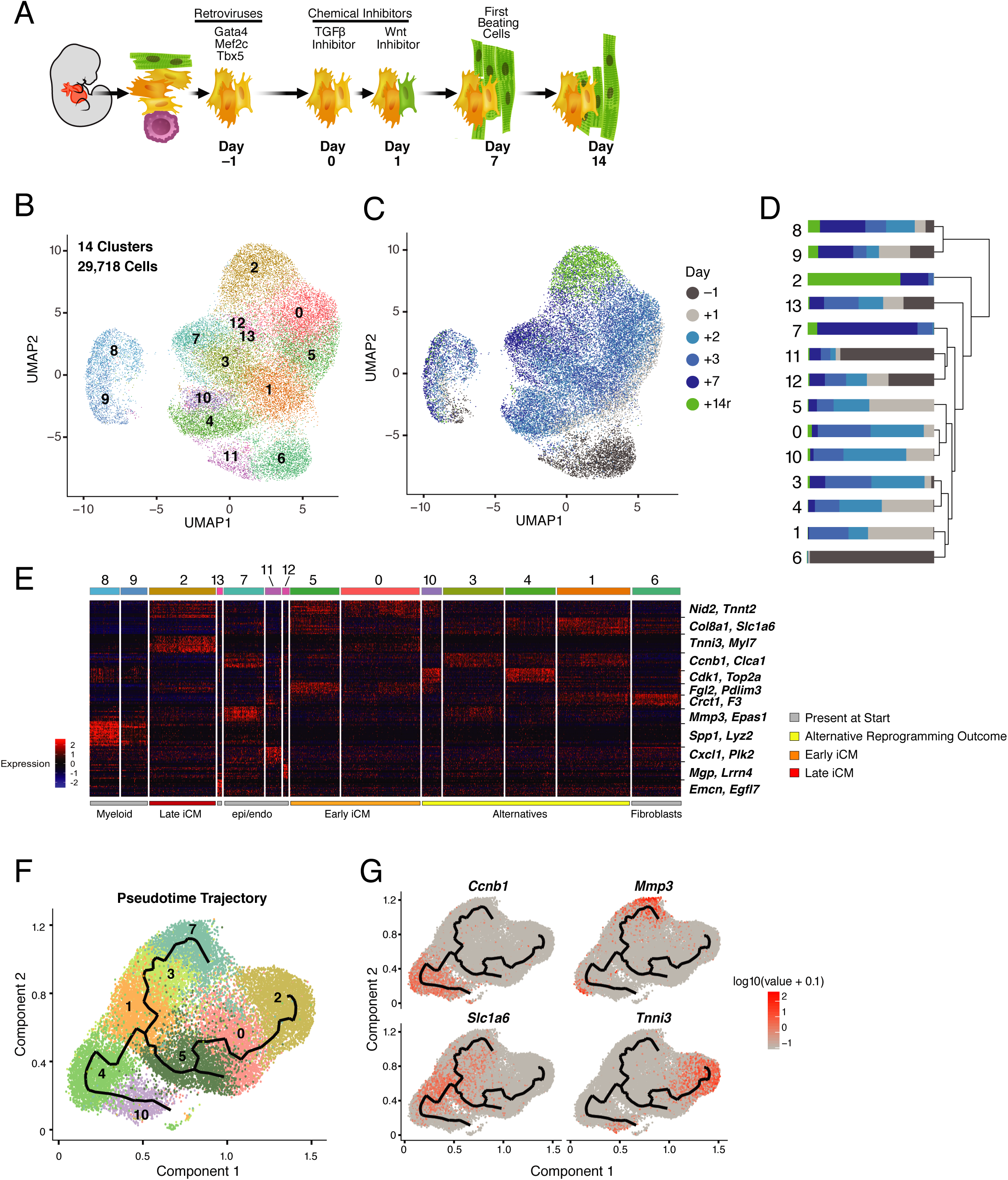
Single-cell Expression Analysis of Direct Cardiac Reprogramming. (A) Schematic of reprogramming system. Tissue explants from αMHC-GFP+ neonatal mouse pups are cultured for ∼10 days, allowing expansion of fibroblast population. Thy1+ cells, largely fibroblasts, are isolated by fluorescence-activated cell sorting (FACS), infected with Gata4, Mef2c, and Tbx5 (GMT) retroviruses, and cultured with chemical inhibitors as indicated for up to 14 days as some reprogrammed cells mature into spontaneously beating iCMs. (B) UMAP visualization of 29,718 cells colored by cluster association identified by a graph-based clustering approach. Cluster numbers as indicated. (C) UMAP visualization of cells as in (B), colored by collection time (days) before and after transduction with GMT. 14r indicates cells collected at day 14 and sorted using the αMHC-GFP reporter (“r” denotes reporter-positive sample). (D) Stacked bar plot indicating the relative contribution of cells from each time point (as shown in C) to each cluster (as shown in B). Dendrogram is generated based on a distance matrix constructed in principal component analysis (PCA) space. (E) Heatmap showing expression of top 20 differentially expressed genes for each cluster. Each row shows the fold change in expression of a single gene compared to the population mean of that gene. Red indicates higher expression; blue lower. Order of clusters according to dendrogram relationship depicted in (D). Clusters are identified along the top, with colored bars matching (B). Two marker genes per cluster are labeled at the right. Putative cell identity, determined based on differential expression, is labeled at bottom with colors indicating (grey) cells present in our starting population, (yellow) alternative reprogramming outcome, (orange) early iCMs and (red) late iCMs. (F) Pseudotime trajectory of cells from clusters 0, 1, 2, 3, 4, 5, 6, 7, 10, and 11 generated using Monocle. Cell color is based on cluster color in (B). (G) Expression (log10(UMI+0.1)) of branch marker genes (*Tnni3, Mmp3, Ccnb1,* and *Slc1a6*) visualized in pseudotime trajectory plots.

Transcript information from all samples was aggregated and a graph based clustering approach using principal component analysis and the Louvain algorithm identified 14 distinct transcriptional signatures which were visualized using Uniform Manifold Approximation and Projection (UMAP) (**Figure 1B, Table S1**) (Becht et al., 2018; Butler et al., 2018). Excluding the clusters that exclusively represent day -1 (clusters 6 and 11), all additional clusters identified included cells collected at each time point, highlighting the limited technical variability between our timepoints and the heterogeneous response to GMT (**Figure 1C, D**).

To better understand the biological significance of the 6 main groups of cells identified through hierarchical clustering of our populations, we next identified representative gene signatures for each cluster (**Figure 1E).** A differential expression test revealed 7,395 genes were differentially expressed (p < 0.01, average log fold change > 0.3) across our time course (**Table S1)**. Three populations represented non-fibroblast cell types present in the starting population that were detected throughout our time course: epicardium-derived (clusters 11 and 12), endocardium (cluster 13), and macrophages (clusters 8 and 9) (**Figure 1D-E**, **Figure S1B**). Endocardial cells were identified by expression of *Emcn* and *Egfl7* (cluster 13, **Figure 1E**) (Cavallero et al., 2015). Epicardial cells were detected based on expression of genes such as *Lrrn4* and *Mgp* (cluster 12, **Figure 1E**) (Xiao et al., 2018). Notably, the expression of these genes exclusively within endocardium and epicardium cells throughout our time course does not support a previous report that suggested these genes are expressed at early stages in reprogramming cells but subsequently repressed by GMT expression (**Figure 1C-D**) (Liu et al., 2017). Instead, our analysis suggests these distinct cell types persist in the population in low numbers and were not detected in the previous study due to a limitation in the number of cells captured.

Four additional signatures were identified that putatively represent various stages or outcomes of cardiac reprogramming. The initial stages of reprogramming (early iCMs) are represented by clusters 0 and 5, identified by activation of genes such as *Nid2* and *Tnnt2* and incomplete repression of fibroblast-associated genes such as *Col8a1* and *Ptgs2* (**Figure 1E**, **Figure S1B**). This signature was present within 48 hours of GMT transduction, in agreement with previous reports (Liu et al., 2017; Sauls et al., 2018). Cluster 2 represents late iCMs that express cardiomyocyte-related genes such as *Nppa, Tnni3, Sln,* and *Myl7,* and have downregulated the fibroblast gene program (**Figure 1E**, **Figure S1B**). Unexpectedly, this cluster contained cells collected from days 3 (5% of day 3 cells) and 7 (26% of day 7 cells), as well as reporter-positive cells collected on day 14 (“+14r”; **Figure 1C**), suggesting a reprogrammed state can be acquired rapidly (**Figure 1D**).

The two additional signatures putatively contain alternative reprogramming outcomes. The first exhibits expression of various genes associated with the cell cycle (clusters 4 and 10) such as *Cdk1* and *Top2a* (**Figure 1E**). Alternatively, cells in cluster 1 are more similar to the starting population, but have activated a subset of cell-cycle genes such as *Ccnb1* (**Figure 1E**, **Figure S1B**). These cells are similar to cluster 3 in both UMAP space and based on hierarchical clustering, which has activated genes that become more robustly expressed in cluster 7, such as *Mmp3* and *Ccl7* (**Figure 1C-E**, **Figure S1B**). Cluster 7 is similar to clusters 11 and 12, which are found in the starting population, suggesting cluster 7 represents cells that do not acquire a cardiac fate nor enter the cell cycle (**Figure 1D-E**). Gene ontology (GO) analysis of genes activated in cluster 7 (compared to cluster 11, n=488 genes, average log fold change > 0.3, p < 1×10^-10^) revealed biological processes associated with vasculature and blood vessel development (p = 1.76×10^-13^ and 5.32×10^-13^, respectively). These cells uniquely activate genes associated with vascular developmental processes such as *Epas1, Figf, and Sox9* (**Figure 1E, Table S1**) (Achen et al., 1998; Lincoln et al., 2007; Tian et al., 1997). They also continue to express genes associated with the starting fibroblast state, such as *Dcn* and *Tbx20* (**Table S1**). Therefore, unlike direct neural reprogramming, we do not detect the emergence of an alternative cell type in our experiments (Treutlein et al., 2016).

To better understand the associations between identified clusters and establish a transcriptional trajectory of the reprogramming process, we next ordered clusters in pseudotime using Monocle (Cao et al., 2019). Clusters containing alternative cell types identified in the starting population were eliminated (clusters 8, 9, 11, 12, and 13) from this analysis as they do not reflect the process or outcome of reprogramming. The starting fibroblasts (cluster 6) were also eliminated as they are too dissimilar to the reprogramming cells, and their inclusion masks the additional dynamics. This analysis indeed revealed a tree structure with three main branches indicating three possible outcomes (**Figure 1F**). One branch reflects an alternative outcome exhibited by progressive activation of *Mmp3*, while activation of cardiac genes such as *Tnni3* and positive markers of cell cycle progression (e.g. *Ccnb1*) occur along separate, distinct trajectories (**Figure 1G**). The inverse relationship between cardiac reprogramming and proliferation is aligned with fibroblast reprogramming towards a pluripotent state which is also negatively impacted by continued proliferation (Xu et al., 2013). Collectively, these data suggest that a reprogramming trajectory can be acquired within 48 hours of GMT transduction but that reprogramming progress occurs at variable rates in individual cells.

### Reprogramming Trajectory is Entered Quickly and Driven by GMT Transduction

To understand how ectopic GMT expression may dictate the observed transcriptional trajectories, we assayed expression of Gata4, Mef2c, or Tbx5 in each cell by generating 5’ single cell RNA sequencing data for 2,593 cells collected on day 1 of reprogramming. This approach circumvented the limitation of our initial analysis whereby the individual ectopic retroviral plasmids could not be distinguished because they each encode the same 3’ polyadenylation sequence that was used to generate cDNA during sequencing library construction. After eliminating the myeloid lineage, we identified 12 clusters within this population, confirming the prompt rate in which cells alter their transcriptional landscape in our system (**Figure 2A**). A pseudotime analysis again identified three main branches in the main trajectory as well as a separate group of clusters (clusters 6, 7, 11, and 12) that were unlinked from the main trajectory (**Figure 2B**).

**Figure 2.**
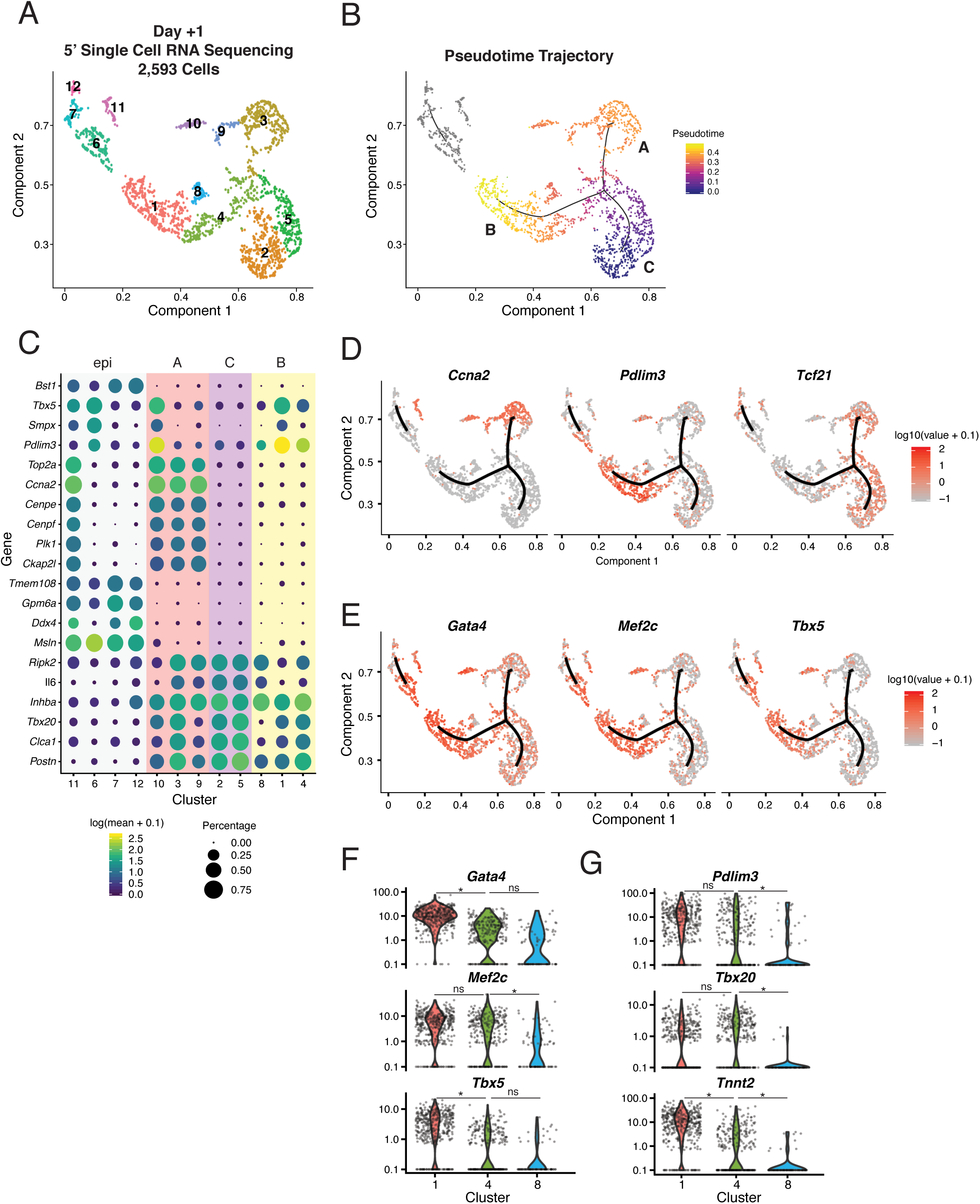
Cardiac Reprogramming Trajectory Is Entered Rapidly. (A) UMAP visualization of 2,593 cells collected at day +1 and colored by Louvain-identified clusters. (B) Pseudotime trajectory identified by Monocle of all cells in (A). Color indicates progression in pseudotime space. Grey indicates disjointed trajectory. Letters indicate main branches in the main trajectory. (C) Dot plot showing expression of top marker genes from each cluster based on specificity calculated by Moran’s I test. The color of the dot represents the average expression level (log10(UMI+0.1)) while the size of the dot represents the percentage of cells expressing the gene. Background color indicates branches from (B). (D) Expression (log10(UMI+0.1)) of branch markers *Ccna2, Pdlim3,* and *Tcf21* visualized in a UMAP trajectory plot. (E) Expression (log10(UMI+0.1)) of *Gata4*, *Mef2c,* and *Tbx5* visualized in a UMAP trajectory plot. (F) Violin plots depicting normalized UMI levels for *Gata4, Mef2c, and Tbx5* in cluster 1, 4, and 8, colored based on cluster identification depicted in (A). Stars indicate negative binomial adjusted p-values. (G) Violin plots depicting normalized UMI levels for *Pdlim3, Tbx20,* and *Tnnt2* in clusters 1, 4 and 8, colored based on cluster identification depicted in (A). Stars indicate negative binomial adjusted p-values.

The three branches identified in the main trajectory represent gene signatures analogous to those presented in Figure 1 based on a differential expression analysis; cells that are likely proliferating (A, 22%), reprogramming cells (B, 33%), and fibroblast-like cells (C, 29%) (**Figure 2C**, **Table S2**). Evaluation of GMT expression revealed their collective expression is observed in branch B, which contains cells that have activated markers of a cardiomyocyte fate (e.g. *Pdlim3*, *Nid2*, *Smpx,* and *Tnnt2)* and downregulated genes associated with a fibroblast identity (*Tcf21, Postn,* and *Tbx20*) (**Figure 2C-E**, **Table S2**). *Gata4* is expressed in cardiac fibroblasts and is therefore detected throughout the population; however, it is increased 1.6 fold in cluster 1 compared to cluster 2. GMT is also expressed in cluster 10, which lies within branch A, suggesting expression of GMT can lead to a partially reprogrammed state even if the cells enter a proliferative state (**Figure 2C-E**). While this type of reprogramming may not produce more advanced iCMs, it suggests proliferation does not prevent the initial stages of reprogramming (Liu et al., 2017). In contrast to branches A and B, the fibroblast-like cells that populate branch C exhibited only baseline levels of all three factors (**Figure 2E**). This analysis confirms that this cell type arises from fibroblasts that do not express ectopic GMT, rather than representing a newly acquired state driven by transduction with one or two factors.

GMT expression was also detected within the unlinked trajectory in clusters 6 and 11 (**Figure 2E**). These clusters contain epicardial cells expressing *Ddx4, Lrrn4, and Msln* (**Figure 2E**, **S2A, Table S2**). Cluster 6 additionally upregulated early markers of the iCM trajectory such as *Pdlim3*, *Smpx,* and *Cd24a*, suggesting that, although unlinked from the main trajectory, this cell type may be capable of acquiring a cardiomyocyte-like gene expression signature upon transduction with GMT (**Figure 2C-D, Table S2**). A similar cell cycle-related phenomenon was also observed in the epicardial cells, as cells in cluster 11 entered the cell cycle and activated expression of *Pdlim3* and *Smpx* (**Figure 2C-E, S2B Table S2**).

To identify variables that may dictate progress in the reprogramming trajectory, we next compared the gene expression profiles of clusters 1 and 4 as they represent early, yet distinct iCM reprogramming states. Examination of GMT expression levels found a statistically significant difference in *Gata4* (negative binomial adjusted p-value = 9.58×10^-45^) and *Tbx5* (negative binomial adjusted p-value = 2.08×10^-15^) between clusters 1 and 4, but not *Mef2c* (**Figure 2F, S2B**). Cluster 1 exhibits stronger upregulation of early markers of reprogramming (e.g. *Smpx, Tnnt2, and Cd24a*) and downregulation of fibroblast-associated genes (e.g. *Postn, Tbx20,* and *Sdpr*) (**Figure 2C, F-G, S2B, Table S2**). Therefore, while robust expression of *Mef2c* and *Tbx5* is required, this variation suggests lower levels of *Gata4* may limit the rate of reprogramming but still allow initiation of the process. While cluster 8 is most similar to clusters 1 and 4, it has not activated markers of reprogramming but it has downregulated markers of the starting fibroblasts (**Figure 2F-G**). There was a significant difference in *Mef2c* expression (negative binomial adjusted p-value = 1.17×10^-08^), further supporting the necessity of robust expression of this gene. Ectopic expression of GMT could be limited by post-transcriptional mechanisms that influence protein stability or a subset of cells may have received fewer viral copies.

### Chromatin Remodeling Occurs within 72 Hours of GMT Expression

To identify the dynamics in chromatin accessibility underlying the aforementioned transcriptional changes, we performed ATAC-seq on αMHC-GFP positive cells collected at five time points during reprogramming (days 2, 3, 7, 14, and 21), and compared regions of accessible chromatin to those detected in the starting fibroblast population. This analysis identified 100,691 total dynamic regions, which included a rapid gain of accessibility by day 2 of reprogramming at the early reprogramming marker gene *Slc6a6* and cardiac *Tnnt2* loci (**Figure 3A**, **Figure S3A-B, Table S3**). Principal component analysis of the genome-wide chromatin accessibility data shows extensive chromatin remodeling by day 2, in agreement with the transcriptional dynamics presented in Figure 1 **(Figure S3C)**.

**Figure 3.**
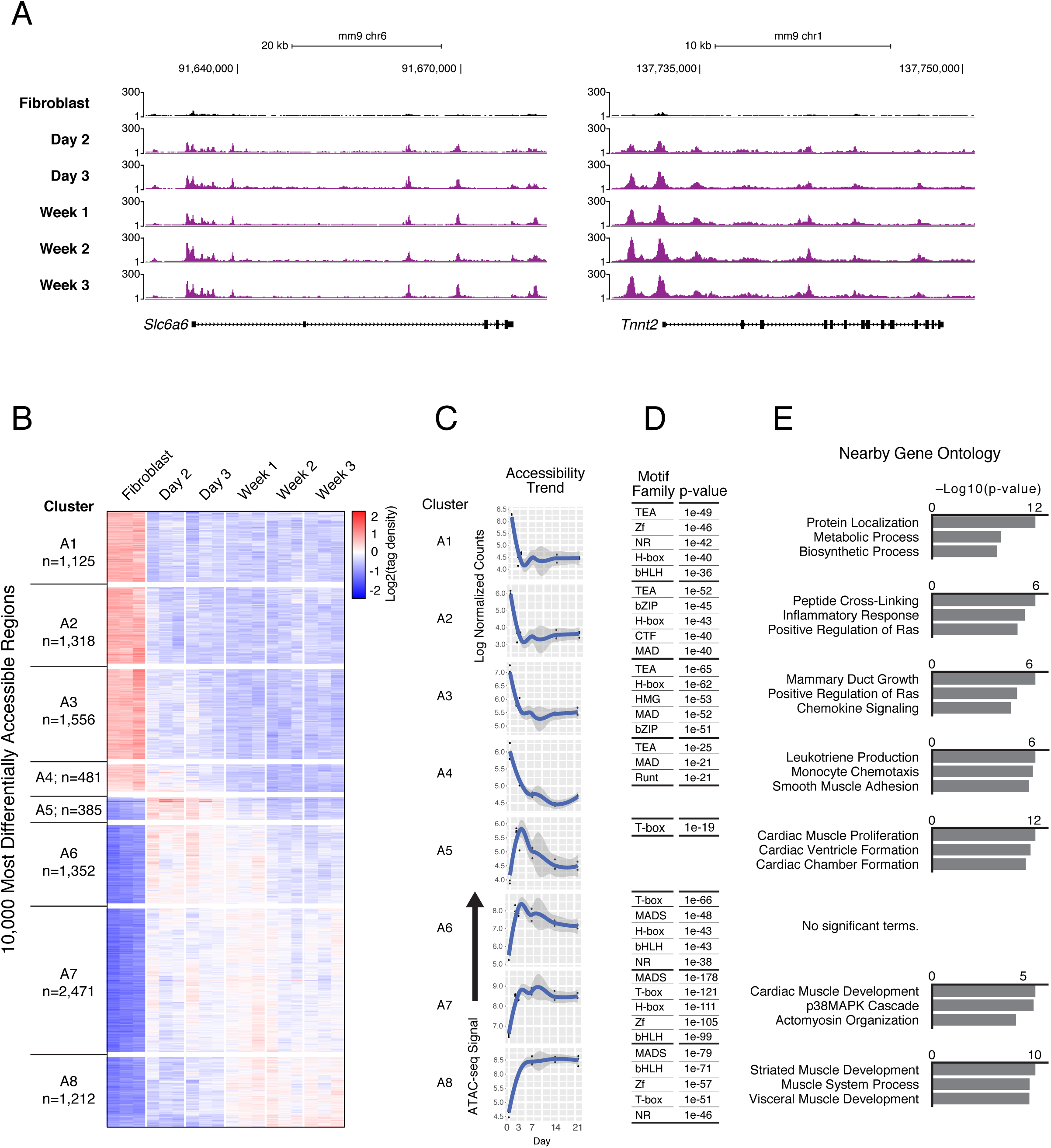
Reprogramming Initiates Widespread Chromatin Accessibility Changes. (A) Profiles display gain of ATAC-seq signal in iCMs harvested at day 2, day 3, week 1, week 2, and week 3 after reprogramming induction with GMT (purple) compared to fibroblasts (black) near the early reprogramming marker gene *Slc6a6* (left) and cardiomyocyte gene locus *Tnnt2* (right). Only non-duplicate read pairs mapping within 500bp were retained, and track height representing the density of reads from merged biological replicates (n=3) was normalized to sample read depth of the condition with the highest number of mapped read pairs. (B) Heatmap displays hierarchical clustering of 10,000 regions with most differentially accessible chromatin status in αMHC-GFP+ iCMs harvested at day 2, day 3, week 1, week 2, and week 3 after reprogramming induction with GMT, compared to fibroblasts infected with dsRed control retrovirus. Clustering highlights 8 distinct patterns (A1-A8). Red indicates accessibility gain, while blue indicates decrease. The data plotted are row-normalized log2 counts for these 10,000 regions across all samples, and the number of differentially accessible regions per cluster is depicted by n. (C) Medoid plots show ATAC-seq tag density over time at dynamic regions from each cluster A1-A8 that are representative of overall trends. Grey region represents the 95% confidence interval of the standard error of the mean signal profile of the representative region. (D) Tables list transcription factor families with motifs significantly enriched (p ≤ 1e-19) within dynamic regions from each cluster compared to all stably accessible (non-dynamic) regions. P-values listed are from the top ranked motif from each transcription factor family. Complete motif enrichment results are available in Table S4. (E) Bar charts show top three ranked biological process terms enriched in dynamic regions from each cluster compared to all stably accessible (non-dynamic) regions. Enriched annotations were determined using GREAT (McLean et al., 2010).

To uncover factors that direct the most robust changes in chromatin accessibility, we performed principal component analysis and hierarchical clustering on the 10,000 most differentially accessible regions identified during our time course and found eight primary patterns (**Figure 3B**). Approximately half of the most dynamic regions (n=4,480) identified in our time course lost accessibility during differentiation, while the other half gained accessibility (n=5,520). Regardless of chromatin remodeling dynamics, the vast majority of changes occurred distal from transcriptional start sites, with dynamic regions underrepresented in promoter proximal regions (p = 2.2e-16; **Figure S3D**). The majority of regions that lost accessibility exhibited this change within 3 days of GMT induction (**Figure 3B, C; clusters 1-4**). Motif enrichment analysis identified the TEAD family as most associated with loss of chromatin accessibility (**Figure 3D, Table S4**). The TEA transcription factors *Tead1* and *Tead4* are expressed throughout reprogramming (**Table S5, Figure S3E**) and promote cell growth and proliferation through interactions with Hippo pathway components (Yoshida, 2008). Nuclear exclusion of TEAD proteins occurs during cell cycle exit (Ota and Sasaki, 2008).

In contrast to the similar dynamics observed in clusters that exhibited the strongest loss of accessibility, there were multiple distinct patterns associated with gain of accessibility. Cluster A5 demonstrated a gain in chromatin accessibility by day 2, but then exhibited a return towards the fibroblast accessibility state at later time points, suggesting accessible chromatin at those sites was not stabilized (n=385, **Figure 3B, C**). This cluster showed limited enrichment of transcription factor sequence motifs, which may have prevented stable GMT binding similar to findings reported for Mef2c, Gata4 and FOXA2 whereby a lack of motif leads to transient sampling rather than stable binding (**Figure 3D, Table S4**) (Andzelm et al., 2015; Donaghey et al., 2018). Cluster 6 demonstrated a similar initial trend as cluster 5; however, the extent of the loss at later time points was reduced (n=1352, **Figure 3B-C**). Clusters 7 and 8 represent the majority of regions associated with a gain in accessibility, and the maximum gain in these regions was observed after day 3 (n=2471, 1212, **Figure 3B-C**). Regions in these clusters maintained higher levels of accessibility over the time course, compared to clusters 5 and 6, and also contained significant enrichment of multiple motif families (**Figure 3D, Table S4**).

To assess the potential functional roles of each cluster, we annotated regions exhibiting changes in chromatin accessibility using GREAT (**Figure 3E, Table S6**) (McLean et al., 2010). Regions that exhibited loss of accessibility were associated with the inflammatory response (cluster 2) and monocytes (cluster 3), supporting a previous report that ZNF281 promotes reprogramming through repression of inflammatory signaling pathways (Zhou et al., 2017). Clusters 7 and 8, which gain and maintain accessibility, were associated with cardiovascular terms such as cardiac and striated muscle development. Regions that did not maintain accessibility in cluster 5 were also associated with cardiac function (**Figure 3E, Table S6**). It remains possible that while Tbx5 may transiently sample those sites during reprogramming, it requires developmentally regulated binding partners to initiate and/or stabilize the interactions with DNA that are not robustly expressed in our system, such as Eomes (McLane et al., 2013). In summary, based on the motifs revealed by the ATAC-seq data, we can deduce the identity of additional transcription factors that may be responsible for promoting the cardiac state by increasing accessibility at cardiac loci and/or inhibiting other, alternative cell types throughout the reprogramming process.

### Mef2c and Tbx5 Binding Is Associated with Changes in Chromatin Accessibility

To understand how DNA binding by GMT underlies the early changes in chromatin accessibility, we next performed ChIP-seq at day 2 of reprogramming to assess Gata4, Mef2c, and Tbx5 occupancy. Unlike the ATAC-seq presented in Figure 3, this experiment was performed in immortalized neonatal cardiac fibroblasts to obtain sufficient numbers of cells; therefore, we created a new, matched ATAC-seq dataset for integration with the ChIP-seq data. This analysis identified 5,100, 6,904, and 5,307 peaks for Gata4, Mef2c, and Tbx5, respectively (**Figure 4A, 4B, S4A**). Assignment of peaks to the nearest TSS showed that the majority of Gata4 and Mef2c peaks were located further than 2kb from the nearest TSS, while 45% of Tbx5 peaks were within 2kb of a TSS, suggesting that Tbx5 may have the most direct influence on changes in gene expression (**Figure S4B**).

**Figure 4.**
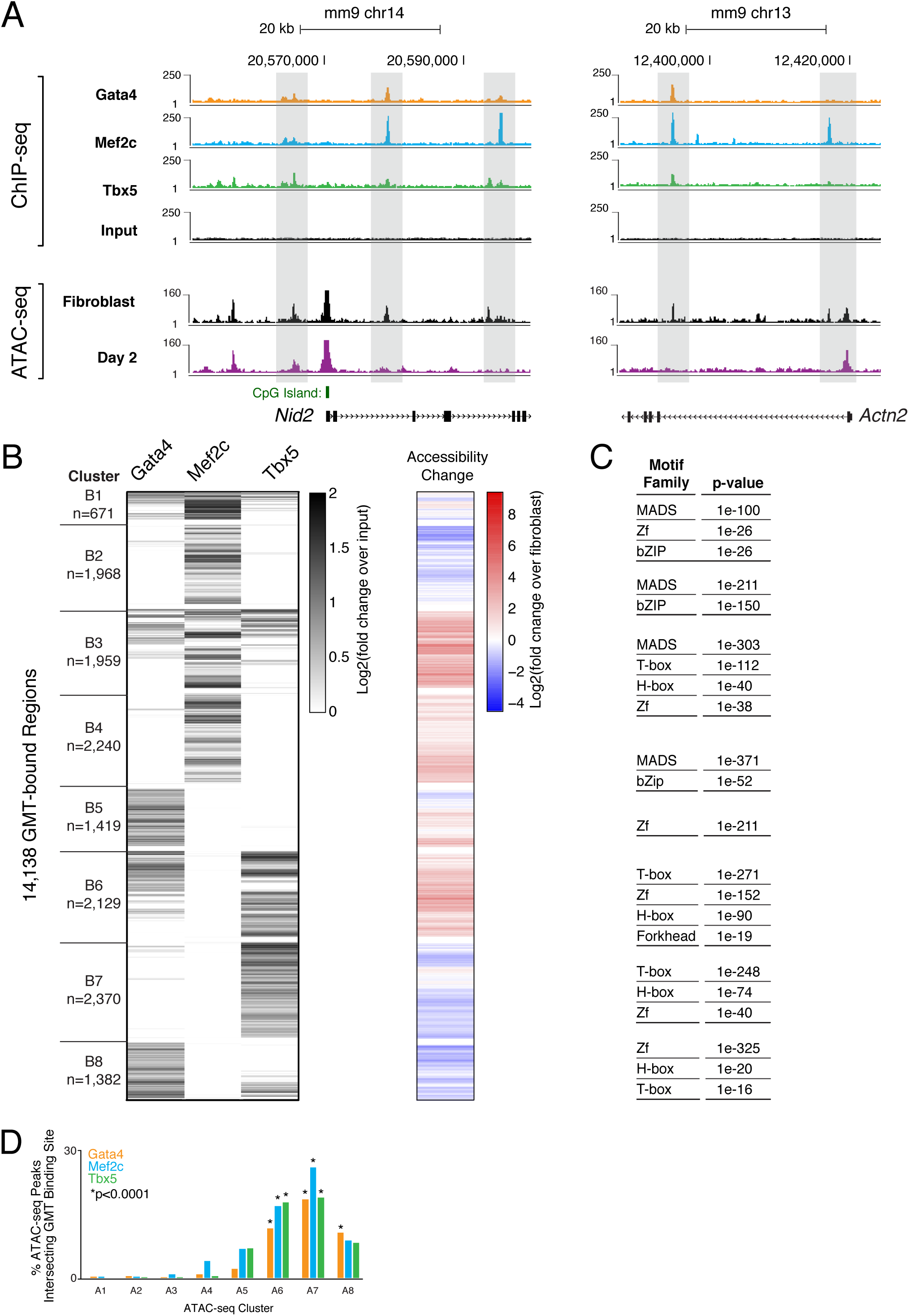
Chromatin Accessibility Dynamics Associated with Gata4, Mef2c, and Tbx5 Occupancy during Reprogramming. (A) ChIP-seq profiles display GMT binding in unsorted cells at day 2 of reprogramming near the early reprogramming marker gene locus *Nid2* (left panel) and the cardiomyocyte locus *Actn2* (right panel) at day 2 of reprogramming. Grey rectangles indicate regions bound by Gata4 (orange track), Mef2c (blue track), and/or Tbx5 (green track). ATAC-seq signal from unsorted cells at day 2 of reprogramming (purple track) and fibroblasts infected with control dsRed retrovirus (black track). ChIP-seq track height is normalized to sample read depth of condition with highest number of reads. ATAC-seq track height is normalized to sample read depth of condition with highest number of mapped read pairs. Tracks are representative of read density from merged biological replicates (Gata4 ChIP, n=6; Mef2c ChIP, n=4; Tbx5 ChIP, n=4; for all ATAC-seq conditions n=3). (B) Heatmap displays hierarchical clustering of ChIP-seq peaks. Grey scale displays average tag density across replicates, normalized to input, with black indicating an increase in tag density in sample over input. At right, co-clustered changes in accessibility. Blue-red color scale indicates tag density of ATAC-seq at day 2 of reprogramming, normalized to fibroblast. Red indicates a relative increase in accessibility, while blue indicates a decrease. Co-clustering reveals 8 distinct patterns (B1-B8). The counts are normalized for differences in sequencing depth between samples using upper quartile normalization separately for the the ChIP and the input samples of each transcription factor. Data for individual replicates including inputs are shown in **Figure S4A**. (C) Tables display transcription factor families with motifs significantly enriched within GMT-bound regions from each cluster, compared to stably accessible (non-dynamic) regions. P-values listed are from the top ranked motif within each transcription factor family, calculated using the cumulative hypergeometric distributions. (Full enrichment results available in **Table S7**.) (D) Bar plot displays the percentages of accessible chromatin regions (ATAC-seq peaks) from each cluster (A1-A8 in Figure 3B), bound by Gata4, Mef2c, or Tbx5. Statistical significance determined using Fisher exact test for overrepresentation of overlapping ChIP-seq peaks in one cluster compared to all dynamic region clusters.

To reveal relationships between reprogramming factor binding and chromatin accessibility, we performed hierarchical clustering on the merged region set (n=14,138) bound by Gata4, Mef2c, and/or Tbx5 during reprogramming, including changes in chromatin accessibility, and identified eight primary patterns (**Figure 4B, S4A**). Only two groups (clusters B1 and B3, n=2,630) included binding of all three factors. However, these clusters exhibited disparate chromatin dynamics. Cluster 1 contained regions that were accessible in the starting population, and their accessibility increased slightly by day 2 (**Figure 4B**). Regions within cluster 3 were alternatively inaccessible in the starting fibroblasts and their accessibility increased (6.33-fold mean increase by day 2) (**Figure 4B**).

This intersection revealed two clusters containing regions bound by Mef2c alone (clusters B2 and B4; n=1,968 and n=2,240, respectively), each displaying opposing trends in chromatin accessibility **(Figure 4B)**. While regions in cluster 2 lost accessibility on average, regions in cluster 4 experienced a 2.55-fold mean increase, which may be linked to the greater representation of a Mef2 motif in cluster B4 vs cluster B2 (61.17% vs 49.44%, respectively) (**Figure 4C**, **Table S7**). While the trend identified in cluster 4 suggests the machinery Mef2c requires to promote chromatin remodeling is active during reprogramming, we did not observe significant chromatin remodeling at regions bound by Tbx5 alone (cluster B7, n= 2,370) or Gata4 alone (cluster B5, n=1,419) (**Figure 4B**). Cluster B7 exhibited little change in accessibility during reprogramming (0.64-fold mean change), while regions bound by Gata4 and Tbx5 together (cluster B6; n=2,129), exhibited a 3.84-fold mean increase in accessibility, suggesting synergistic binding of Gata4 and Tbx5 has a positive impact on chromatin remodeling at those regions (**Figure 4B**).

We next identified potential non-GMT cofactors within these regions by searching for known motifs enriched within ChIP-seq peaks, and summarized them based on TF family (**Figure 4C, Table S7**). As expected, top-ranked motifs corresponded to families of reprogramming transcription factors; however, additional motif families were also significantly enriched, including bZip, Homeobox, and Forkhead (**Figure 4C, Table S7**). These families include transcription factors such as Atf1/2/3/7, Fosl2, Jun, and Bach2 (bZip); Tgif1/2 and Meis1 (H-box); and Foxm1 (Forkhead), all of which are expressed during reprogramming (**Figure S3E**).

Finally, we analyzed the binding of Gata4, Mef2c, and Tbx5 at regions that exhibited the most dynamic chromatin accessibility changes during reprogramming (regions from **Figure 3B-C**). We detected no enrichment of GMT binding at day 2 in regions that lose accessibility during reprogramming (clusters A1-4), while we detected statistically significant enrichment of GMT binding at day 2 in almost all regions that gained accessibility during reprogramming (clusters A6, A7, and A8) (**Figure 4D**). Cluster A5 is the only ATAC-seq cluster that gained accessibility at day 2 of reprogramming without significant enrichment of binding by reprogramming factors, providing a potential explanation for why this cluster exhibits only a transient accessibility gain (**Figure 4D, 3B-C**). Taken together, this suggests that chromatin accessibility dynamics directed by binding GMT are context-specific.

### Transcription Factor DNA Occupancy Defines Chromatin Accessibility Trends

To understand how GMT binding, individually and in combination, is related to changes in chromatin accessibility, we next performed ChIP-seq and ATAC-seq at day 2 with single factors (SF) ectopically expressed, and additionally, with pairs of factors ectopically expressed (DF) (**Figure 5A**). Principal component analysis reveals that each individual factor’s binding pattern differed from the pattern detected during reprogramming when all three factors were present (**Figure 5B**). A large shift occurred for Gata4, whose binding became similar to Tbx5 during reprogramming with all factors, supporting a previously reported cooperative binding relationship between these two transcription factors in developing mouse and human cardiomyocytes (Ang et al., 2016; Luna-Zurita et al., 2016; Maitra et al., 2009). Mef2c exhibited a decidedly distinct binding pattern compared to Tbx5 and Gata4, but was altered by the addition of Gata4 and Tbx5 (**Figure 5B).**

**Figure 5.**
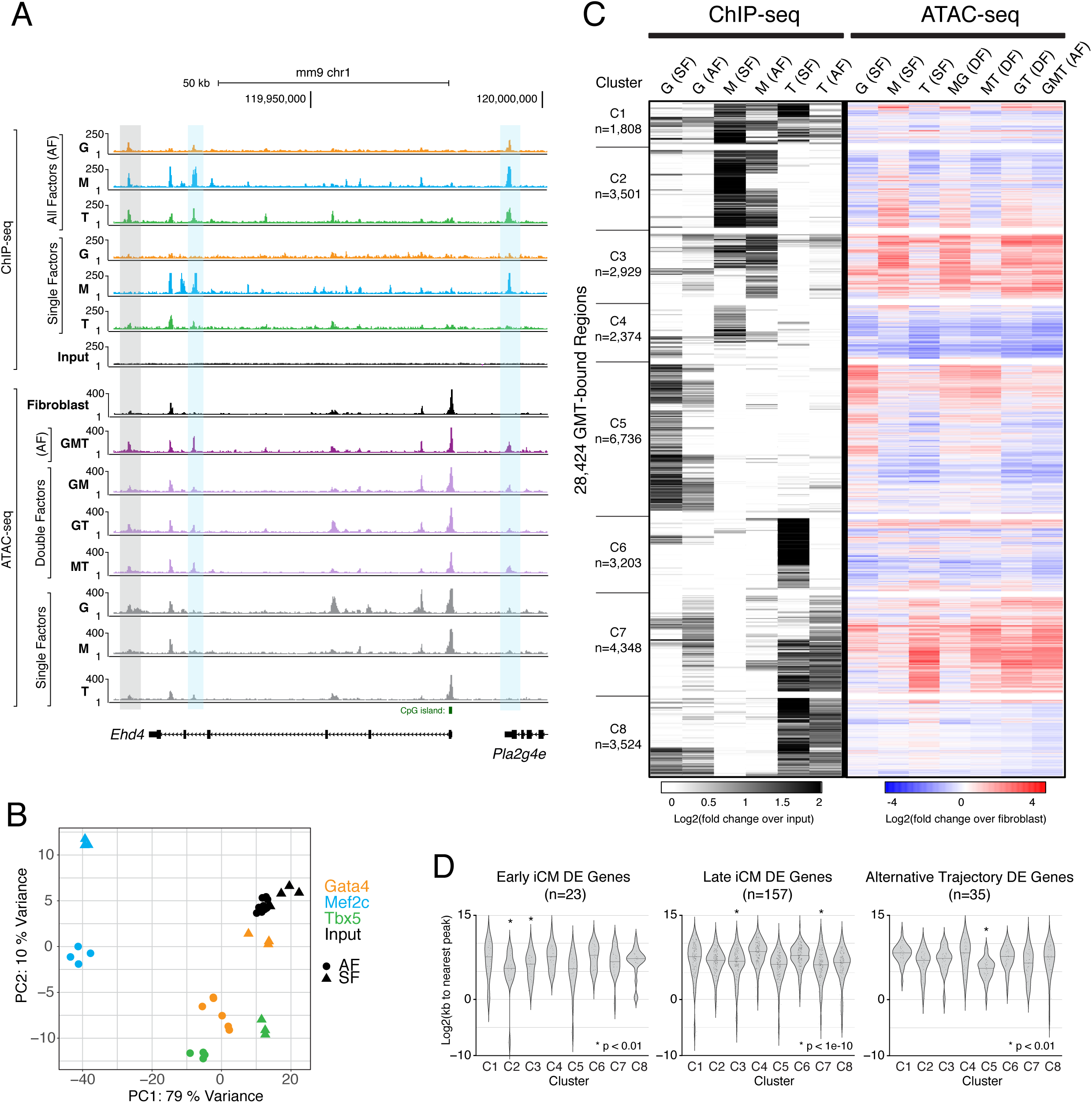
Chromatin Accessibility Dynamics Associated with Gata4, Mef2c, and Tbx5 Occupancy Following Independent and Combinatorial Expression. (A) Profiles display ChIP-seq signal for GMT in single factor (SF) and all factor (AF) conditions, and ATAC-seq signal in SF, double factor (DF), and AF conditions, near the early reprogramming marker gene locus, *Ehd4*. Grey rectangle highlights region bound by Gata4 (orange tracks), Mef2c (blue tracks), and Tbx5 (green tracks) in AF condition, without binding by any of these factors in SF conditions. Blue rectangles highlight regions bound primarily by Mef2c when ectopically expressed alone (SF), but bound by Gata4, Mef2c, and Tbx5 when all factors are expressed (AF). Profiles represent read density from merged biological replicates normalized to read depth (Gata4 AF ChIP, n=6; Mef2c and Tbx5 AF ChIP, n=4; SF ChIP conditions, n=3; all ATAC-seq conditions, n=3). (B) Principal component analysis of all ChIP-seq replicates based on 500 most variable regions among all conditions. Gata4 ChIP-seq samples are shown in orange, Mef2c in blue, Tbx5 in green, and input in black. Single factor (SF) expression conditions indicated by a triangle (▴) and all factor (AF) expression indicated by a circle (●). (C) Heatmap displays hierarchical clustering of ChIP-seq peaks, with grey scale displaying average tag density across replicates, normalized to input (left). Black indicates an increase in tag density in sample over input. At the right, co-clustered tag density of ATAC-seq, normalized to fibroblast. Red indicates a relative increase in accessibility, while blue indicates a decrease. Co-clustering reveals 8 distinct patterns (C1-C8). Conditions include exogenously expressed single factors (SF), double factors (DF), and all reprogramming factors (AF), with unsorted cells harvested at day 2 of reprogramming. (D) Violin plots show distribution of distances from ChIP-seq peaks in each cluster to nearest TSS of differentially expressed genes (DE genes) in cells from “Early iCM”, “Late iCM”, and “Alternative Trajectory” scRNA-seq clusters. Internal lines indicate median values. Wilcoxon p-values were determined per cluster, by comparing distances to TSSs of DE genes with distances to all TSSs, and adjusted for multiple testing using Bonferroni correction.

Hierarchical clustering of the merged 28,424 region set bound by Gata4, Mef2c, and/or Tbx5 when all factors (AF) or single factors (SF) were ectopically expressed, together with changes in chromatin accessibility, resulted in 8 distinct clusters (**Figure 5C)**. Cluster C1 represents regions bound by all three factors, regardless of whether the other reprogramming factors are present; however, most clusters were driven by binding of a single factor (**Figure 5C**). Clusters C2, C4, C5, and C6 represent clusters bound in single factor conditions that are refined by addition of all factors (**Figure 5C, S5A).** Binding of a single factor in C2 and C5 is associated with a concomitant increase in chromatin accessibility. These single binding events refined by the addition of the other reprogramming factors coincide with overrepresentation of the single factor’s sequence motif (**Figure S5B, Table S8**). Regions in C5 exhibit an increase in chromatin accessibility only when Gata4 is present alone, which may suggest these regions act as regulatory elements in alternative cell types for which Gata4 is involved but Tbx5 and Mef2c are not, such as at the TSS of the endothelial gene *Lecam2* (**Figure S5A**). Similarly, Mef2c was sufficient to induce an increase in chromatin accessibility in a subset of regions within C2 that is abrogated in the reprogramming condition (mean fold changes, M= 1.75, MG=1.24, MT=1.12, AF=0.84) (**Figure 5C; Table S9**). Mef2c and Tbx5 single binding events were also associated with a loss of chromatin accessibility (C4 and C6), suggesting their ability to interact with DNA and alter chromatin accessibility is context dependent. C8 confirms the divergent response to Tbx5 binding as a subset of these regions are also bound by Gata4, and are associated with minimal changes in chromatin accessibility (**Figure 5C**). We did not identify a cluster in which Gata4 is independently capable of increasing chromatin accessibility, suggesting its function in the reprogramming context is downstream of epigenetic remodeling.

We noted unique trends in clusters C3 and C7. They are dominated by regions that exhibit binding of all three factors in the all factor reprogramming condition, but limited binding in the single factor conditions, suggesting cooperation between reprogramming factors facilitates DNA binding at these regions (**Figure 5C**). Such areas demonstrated the greatest average increases in chromatin accessibility among regions included in this analysis. While Mef2c binding was sufficient to promote a 3.09-fold increase in chromatin accessibility in cluster C3, Tbx5 comparably directed a 3.08-fold increase in cluster C7 (**Figure 5C, Table S9**). This suggests that chromatin changes induced by Mef2c and Tbx5 in these clusters create a chromatin landscape amenable to the binding of the additional reprogramming factors, adding another layer of regulatory complexity.

We next ascertained the relationship between these clusters and the transcriptional signatures presented in Figure 1. To that end, we defined genes that uniquely represent early iCMs, late iCMs and untransduced fibroblasts based on our single cell RNA sequencing analysis (p<0.0001), and calculated the distance from the TSS to the closest dynamic region included in the 28,424 region set. Both clusters C2 and C3, which represent regions whose chromatin dynamics are largely associated with Mef2c binding, were significantly associated with genes marking “Early iCM” populations (p < 0.001, **Figure 5D, left**). Clusters C3 and C7 were associated with “Late iCM” marker genes and an increase in occupancy by all three reprogramming factors during reprogramming (p < 1e-10; **Figure 5D, middle**). The association between C3 and C7 with expression of late iCM markers suggests that the “Late iCM” trajectory in particular results from cooperative binding by all three reprogramming factors while Mef2c facilitates the initial gene expression changes that define early iCMs based on the significant association between C2 and early iCM genes.

We next asked if any of the observed chromatin dynamics occurred near genes that represent the alternative trajectory, which contains untransduced cells that do not reprogram. Indeed, we found that regions in cluster C5 that were bound by Gata4 only in the SF condition were significantly closer to genes that represent alternative trajectory (p < 0.001; **Figure 5D, right**). The lack of chromatin accessibility changes in the DF conditions in C5 indicate that neither Mef2c (MG) nor Tbx5 (GT) bind to or prevent Gata4 from binding to or altering chromatin accessibility at these regions, supporting our conclusion that these cells represent an untransduced population that expresses and is regulated by endogenous levels of Gata4 (**Figure 5A, C**). Cumulatively, these data reveal the complexity of the mechanisms through which transcription factors influence each other, both enabling and refining one another’s ability to bind DNA and affect accessibility changes.

### Modeling GMT-Induced Transcriptional Changes

In an effort to discern additional transcription factors that direct the initial stages of reprogramming, we devised a multivariate random-forest based machine learning approach to predict which transcription factor sequence motifs are most associated with transcriptional changes observed at day 2. The first iteration of our model incorporated the motifs of expressed transcription factors that lie within accessible chromatin, and the motif’s distance to the transcription start site (TSS) of a differentially expressed gene (**Figure 6A**). However, we found a low correlation between associations within 2kb or greater than 500kb from a TSS, which subsequently led us to focus on motifs within 2-500kb of a differentiationally expressed gene (Supplemental Methods). This supports a previous report that suggested regulatory events >2kb from the TSS have more influence on gene expression dynamics than more proximal motifs (Pliner et al., 2018). Using our model, we saw an average Pearson correlation value of 0.37 between observed fold-change and the predicted fold-change of gene expression when executing the full model with differential accessibility data from fibroblasts compared with day 2 of reprogramming, and sequence-based motifs from all transcription factors. The correlation was stronger than described in previous reports that utilized ATAC-seq data to infer genomic features involved in gene expression (Dong et al., 2012). Limiting the model to accessible reprogramming factor motifs at day 2 without considering accessibility changes over time led to an average correlation of 0.19, or a correlation of 0.20 for a model using only ChIP-seq validated reprogramming factor motifs accessible at day 2 (Supplemental Methods). Our model predicted 48 motifs significantly associated with transcriptional changes observed during the fibroblast to day 2 reprogramming transition (**Figure 6B**). These data suggest additional transcription factors are indeed involved in regulating gene expression dynamics during reprogramming as most expression dynamics cannot be explained by a limited set of transcription factors.

**Figure 6.**
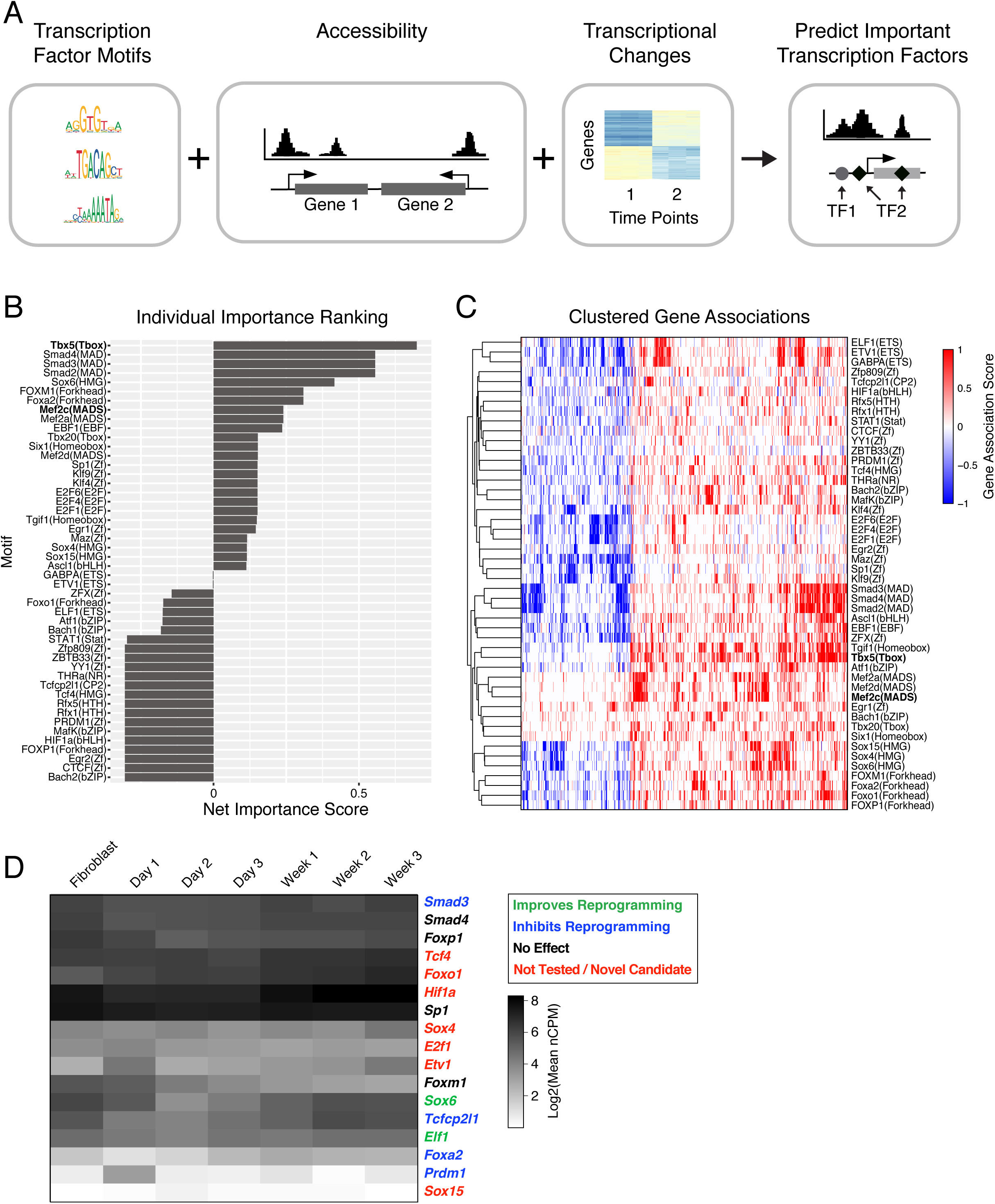
Model of Transcription Factor Association with Changing Rate of Gene Expression over Time. (A) Schematic representation of input features for the multivariate random-forest based machine learning approach, including known transcription factor (TF) motifs, chromatin accessibility, and bulk gene expression, used to predict TFs influencing gene expression changes during reprogramming. TFs are associated with genes using motifs in nearby regions (< 500kb from transcription start site) that change in chromatin accessibility during reprogramming, and the influence of a TF across genes is estimated using targeted maximum likelihood applied to a collection of machine-learning models (Supplemental Methods). (B) Bar chart displays 48 transcription factor motifs significantly associated with gene expression changes observed during the fibroblast to day 2 reprogramming transition, and expressed in at least one time point during reprogramming (Reads Per Kilobase of gene, per Million mapped reads, RPKM > 2). Motif family is listed in parentheses. Transcription factor motifs are ranked by a model-based estimates of motif net influence on fibroblast to day 2 gene expression changes (Bonferroni corrected p-value < 0.05). This net importance score” predicts involvement in transcriptional regulation during reprogramming, with a positive score suggesting an activating influence, a negative score suggesting a repressing influence, and a score close to zero suggesting a combined influence on transcription. (C) Heatmap of clustered gene-association signatures for identified motifs. Rows represent transcription factor motifs; columns are genes associated with these motifs (as described in Supplemental Methods). Red indicates enrichment of the motif nearby a gene, while blue indicates depletion of the motif; white indicates no predicted influence of the transcription factor on the gene. Hierarchical clustering was performed using the Euclidean distance metric as implemented in the pheatmap package in R. (D) Heatmap displays temporal expression levels for top 17 transcription factors predicted to influence reprogramming based on a ranking of putative cooperative relationships between factors. Expression values were generated from bulk RNA-seq of sorted αMHC-GFP+ iCMs collected at each time point in triplicate, and data is shown as log2 mean normalized Counts Per Million mapped reads (nCPM). Transcription factor names listed in green, blue, or black were previously tested during reprogramming induced with Akt1, Gata4, Hand2, Mef2c, and Tbx5 (AGHMT) in adult mouse fibroblasts (Zhou et al., 2017). Of these, factors listed in green significantly improved reprogramming outcome based on αMHC-GFP reporter expression and/or cardiac Tnnt2 expression assayed by antibody staining, while factors in blue inhibited reprogramming by one or both of these metrics. Factors in black had no significant effect on reprogramming outcome. Factors in red have not been tested before, and indicate novel candidate regulators of reprogramming identified in this study. See **Figure S6** for quantification of reprogramming outcomes.

By comparing the rates of expression change between time points in proximity-associated vs non-associated genes for each motif, we ranked the identified motifs based on a “net importance score” (see supplementary methods), which predicts their involvement in the regulation of gene expression during reprogramming (**Figure 6B**). A positive net importance score suggests an activating influence on transcription while a negative score suggests a repressing influence; a score close to zero may indicate a mixed influence. This analysis found that the Tbx5 motif was most associated with gene expression changes during the transition from fibroblast to day 2. Notably, while Mef2 motifs were also among the top ranked set of putative early regulators, Gata motifs were absent from this list, suggesting that Tbx5 and Mef2c are more influential than Gata4 in regulating gene expression changes during the early stages of reprogramming. The lack of Gata4 detection in this prediction combined with our ChIP- and ATAC-sequencing results suggests that it is less impactful than Mef2c or Tbx5 at the earliest stages of reprogramming. Motifs of Smads2/3/4 were also predicted to influence gene expression, supported by previous work from our lab and others that showed TGF**β** inhibition positively influences reprogramming outcome (Ifkovits et al., 2014; Mohamed et al., 2017).

While the random forest model predicted activating and repressive influences for individual transcription factors, it did not reveal if the identified transcription factors target similar or distinct gene sets. We therefore performed hierarchical clustering on the “Gene associating signatures” of each identified motif in order to illustrate putative co-regulatory relationships between multiple transcription factors. (**Figure 6C,** Supplemental Methods). Red indicates motif enrichment is associated with the predicted target gene, while blue indicates a depletion of the accessible motif locations. This results in two major branches, one of which contained many transcription factor motifs associated with cardiomyocyte development such as Mef and Tbx family motifs, as well as Sox, Fox, and SMAD motifs. The Tbx5 motif was most closely linked to changes at regions that also encode Tgif1 motifs, a TGF**β**-induced transcriptional homeodomain-containing repressor (**Figure 6C**).

To predict which of the 48 identified transcription factors may cooperatively impact cardiac reprogramming, we fitted a linear model to determine transcription factor pairs whose influence on gene expression could not be explained by either of the transcription factors alone (i.e., predicted synergistic or antagonistic interaction) (see Supplementary Methods). We ranked the transcription factor motifs by their total number of predicted interactions, and selected 17 top candidate factors that may contribute to reprogramming in a cooperative manner (**Table S10**).

Of our 48 identified transcription factors, 27 have previously been tested as part of a screen for regulators of direct cardiac reprogramming using expression of an αMHC-GFP reporter and/or expression of cardiac Tnnt2 as a measure of reprogramming (Zhou et al., 2017). 41% of the tested factors (11 out of 27) did indeed either enhance or repress reprogramming outcome, compared to the 22% “hit” rate reported from the published screen of 786 genes (transcription factors, cytokines, epigenetic regulators, etc.) pre-selected for their propensity to influence cell fate, thereby validating our machine learning approach. Our selection of factors that may cooperatively impact cardiac reprogramming further improved the ratio of positively tested factors to 60% (6 out of 10) (**Figure 6D, S6**). These results indicate that future studies of the remaining identified candidates may reveal additional factors that can enhance or serve as barriers of cardiac reprogramming.

## DISCUSSION

Here, we interrogated the transcriptional and epigenetic dynamics underlying direct cardiac reprogramming in an *in vitro* mouse cardiac fibroblast system, revealing numerous insights into the mechanisms associated with the cell fate transition from a fibroblast toward a cardiomyocyte. Epigenetic and transcriptional changes occurred broadly within the first 72 hours, and cells destined to reprogram could largely be predicted by virtue of early gene expression dynamics and reprogramming factor expression. Single-cell assays addressed long-standing mysteries regarding heterogeneity and response to combinations of reprogramming factors, clarifying existing interpretation of bulk transcriptome data sets. Integration of GMT DNA occupancy with genome-wide chromatin accessibility in the setting of individual or combinations of transcription factors revealed an interdependency of their binding patterns and suggests a possible mechanism through which they synergize to facilitate successful reprogramming. Finally, a machine learning approach revealed clusters of co-located transcription factor motifs associated with coordinate gene expression changes, pointing to additional factors that may promote or inhibit reprogramming.

Despite similarities to rapid transcriptional and chromatin remodeling seen in other systems, our results highlight differences between direct cardiac reprogramming and other reprogramming types. For example, during the transition from fibroblast to neuron induced using a combination of Ascl1, Brn2 and Mytl (Treutlein et al., 2016), an alternative fate characteristic of skeletal muscle was observed. In contrast, for cells that were successfully transduced with GMT, no major alternative fates were observed. Alternatives may be limited in the cardiac setting, because, as we show here, GMT binding is refined when they are expressed combinatorially, perhaps focusing binding events on cardiac loci. However, Ascl1’s binding is not altered by the addition of other neural reprogramming factors such as Brn2 and Mytl1, suggesting its binding is unrestrained and may occur at regulatory elements employed in the development of multiple cell types (Wapinski et al., 2013). Interestingly, addition of ectopic expression of Ascl1, predicted by our random-forest approach, enhanced cardiac reprogramming suggesting the myogenic outcome in neural reprogramming is not a technical artifact (Zhou et al., 2017).

While reports of direct reprogramming were first documented many years ago, the tools available to dissect the precise molecular mechanism of these processes were limited. Advances in single cell RNA sequencing have created avenues to identify the path a single cell can take to its endpoint, and identify the molecular determinants of these trajectories (Cacchiarelli et al., 2018; Schiebinger et al., 2019). Simultaneous advances in machine learning have improved the information gleaned from scRNA-seq data, as well as the ability to correlate changes between gene expression and chromatin remodeling (Cao et al., 2018; Deng et al., 2019; Lopez et al., 2018; Way and Greene, 2018; Welch et al., 2017). The observation that the vast majority of fibroblasts that expressed GMT proceeded into the induced cardiomyocyte trajectory suggests a higher efficiency among GMT expressing cells than previously recognized. Furthermore, other cell types, such as epicardial cells, have the potential to be partially reprogrammed, although they remain dissimilar to induced cardiomyocytes.

In conclusion, we have developed a comprehensive genomic assessment of transcription factor binding, chromatin state, and transcriptional changes, that reveals the molecular complexity involved in direct cardiac reprogramming. Mechanistic insights provided by integration of multiple datasets have started to reveal how lineage-enriched transcription factors can induce cell fate transitions in a combinatorial fashion, thereby achieving specificity of gene regulation.

## Acknowledgments

We thank members of the Srivastava lab for helpful discussions; J.G. van Bemmel, T.R. Roberts, and K. Claiborn for critical comments and editing on the manuscript; A. Williams, T. Friedrich, and S. Thomas in the Gladstone Bioinformatics Core; Y. Hao and J. McGuire in the Gladstone Genomics Core; M. Cavrois, M. Gesner, and N. Raman in the Gladstone Flow Cytometry Core; S. Elmes in the UCSF Flow Cytometry Core; E. Chow and D. Bogdanoff in the UCSF Center for Advanced Technology Core; and C. Benitez and I. Espineda in the Gladstone Laboratory Animal Resource Center. All protocols concerning animal use were approved by the IACUC at the University of California San Francisco and conducted in strict accordance with the NIH *Guide for the Care and Use of Laboratory Animals.* N.R.S was supported by a National Science Foundation Graduate Research Fellowship, R01 Research Supplement to Promote Diversity in Health-Related Research, and a Ruth L. Kirschstein NRSA Institutional Research Training Grant. C.A.G. is a HHMI fellow of the Damon Runyon Cancer Research Foundation (DRG-2206-14). D.S. is supported by NHLBI/NIH grants (R01 HL057181, U01 HL098179, U01 HL100406), the Roddenberry Foundation, the L.K. Whittier Foundation, and the Younger Family Fund. This work was also supported by NIH/NCRR grant C06 RR018928 to the Gladstone Institutes and NIH P30 AI027763, NIH S10 RR028962 and the James B. Pendleton Charitable Trust to the Gladstone FACS core. All data will be available on GEO upon publication.

## Author Contributions

N.R.S., C.A.G., K.J.B.P., K.S., E.R., A.S., T.M.A.M., and P.Y. performed experiments and generated sequencing libraries. N.S. and R.T. analyzed ATAC- and ChIP-sequencing data. C.A.G. analyzed single cell RNA sequencing data. N.R.S., C.A.G., K.N.I., K.S.P., and D.S. conceived and designed experiments, interpreted the data, and wrote the manuscript.

## METHODS

### Cell Culture

Direct cardiac reprogramming was performed on neonatal mouse cardiac fibroblasts as previously described, using only freshly prepared retroviruses (Ieda et al., 2010; Qian et al., 2013). ChIP-seq and ATAC-seq experiments presented in Figure 5 were performed using an immortalized neonatal cardiac fibroblast cell line, expressing cre-excisable T-antigen, developed by Palmer Yu in the Srivastava lab.

## Sequencing Library Construction

### Single-Cell RNA sequencing (scRNA-seq)

Single-cell RNA-seq libraries were prepared using the Chromium Single Cell 3′ Reagent Kits v2 (PN-120236, PN-120237, PN-120262). Biological replicates were created for four times: minus 1, plus 1, plus 7 and plus 14 days. All libraries were pooled and sequenced using the HiSeq 4000 to a read depth of at least 30,000 reads per cell.

### Bulk RNA Sequencing

Total RNA was isolated using the miRNeasy Micro Kit (Qiagen). Bulk RNA-seq libraries were prepared with ovation RNA-seq system v2 kit (NuGEN). The RNA-seq libraries were analyzed by Bioanalyzer and quantified by QPCR (KAPA). Samples were sequenced at 100PE on the Illumina HiSeq 2500 at the Harvard FAS core.

### Assay for Transposase-Accessible Chromatin with Sequencing (ATAC-seq)

We prepared iCM, single-factor, and fibroblast samples for ATAC-seq as previously described (Buenrostro et al., 2013). Aliquots of 10,000–50,000 cells were spun down (310 RCF for 3 minutes) and washed with 200 μL of chilled PBS. Samples were lysed with 200 μL of chilled lysis buffer (20 mM Tris-HCl (pH 8.0), 85 mM KCl, 0.5% NP-40) and spun down at 500 RCF for 5 minutes. Nuclear pellets were transposed with 25 μL of Tagment DNA Buffer, 2.5 μL of Tagment DNA Enzyme (Nextera Sample prep Kit from Illumina, cat # FC-121-1030), and 22.5 μL of nuclease-free H2O. The samples were incubated at 37°C for 30 minutes and stored at −20°C. Transposed samples were purified using the QIAGEN MinElute Reaction Cleanup Kit (cat #28204). Samples were amplified using 25 μL of NEBNext High Fidelity 2x PCR Master Mix, 1.25 μM Nextera custom primer, 1.25 μM Nextera custom primers with unique barcodes, and nuclease-free H2O. We amplified samples using the following PCR conditions: 72°C for 5 minutes; 98°C for 30 seconds; and cycled at 98°C for 10 seconds, 63°C for 30 seconds and 72°C for 1 minute. Half of each sample was amplified for 12 cycles, MinElute purified and assessed by bioanalyzer for library quality. Samples concentration was quantified by Qbit before pooling. Samples were sequenced at 100PE on the Illumina HiSeq 2500 at either the Harvard FAS core or the UCSF CAT core.

### Chromatin Immunoprecipitation Followed by Sequencing (ChIP-seq)

Cells (10^6^ per ChIP) were crosslinked in 1% formaldehyde in suspension at room temperature for 10 minutes with gentle rotation. Crosslinking was quenched by addition of glycine (final 125 mM), followed by incubation at room temperature for 5 minutes with gentle rotation. Cell pellets were lysed in buffer [20 mM Tris-HCl, pH 8, 85 mM KCl, 0.5% NP-40, protease inhibitors] for 10 minutes on a rotator at 4°C. Nuclei were isolated by centrifugation (2,500 x g, 5 minutes, 4°C), resuspended in nuclei lysis buffer [50 mM Tris-HCl, pH 8, 10 mM EDTA, pH 8, 1% SDS, protease inhibitors] and incubated on a rotator for 30 minutes at 4°C. Chromatin was sheared using a Covaris S2 sonicator for 15 minutes (60-second cycles, 5% duty cycle, 200 cycles/burst, intensity = 5) until DNA was in the 200–700 base-pair range. Chromatin was diluted fivefold in ChIP dilution buffer [0.01% SDS, 1.1% Triton X-100, 1.2 mM EDTA, 16.7 mM Tris-HCl, pH 8, 167 mM

NaCl, protease inhibitors] and incubated with antibody (2 mg/million cells) at 4°C overnight under rotation. Antibodies used are Santa Cruz, Gata4, sc-1237x; Cell Signal Tech, Mef2c 5030; Santa Cruz, Tbx5 (C-20) sc-17866x. Antibody-protein complexes were immunoprecipitated using Pierce Protein A/G magnetic beads at 4°C for 2 hours under rotation. Beads were washed five times (2-minute washes under rotation) with cold RIPA buffer [50 mM HEPES-KOH, pH 7.5, 500 mM LiCl, 1 mM EDTA, 1% NP-40, 0.7% Na-deoxycholate], followed by one wash in cold final wash buffer [1xTE, 50 mM NaCl]. Immunoprecipitated chromatin was eluted at 65°C with agitation for 30 minutes in elution buffer [50 mM Tris-HCl pH 8.0, 10 mM EDTA, 1% SDS]. High-salt buffer [250 mM Tris-HCl, pH 7.5, 32.5 mM EDTA, pH 8, 1.25 M NaCl] and Proteinase K were added and crosslinks were reversed overnight at 65°C. Samples were treated with RNase A and DNA was purified with Agencourt AMPure XP beads. Fragmented ChIP and control (whole-cell extract) DNA was end-repaired, 5′ phosphorylated and dA-tailed with NEBNext Ultra II DNA Library Prep Kit for Illumina (NEB E7645). Samples were ligated to adaptor oligos for multiplex sequencing (NEB E7335), PCR amplified and sequenced on an Illumina NextSeq500.

## Sequencing Data Processing and Analyses

### Single-Cell RNA-Seq Analysis

The 10X Genomics Cell Ranger pipeline was used to demultiplex raw data, align reads, count transcripts and aggregate multiple samples and timepoints. The R packages Seurat v2.3 and Monocle v3 were used for all downstream analyses (Butler et al., 2018; Cao et al., 2019). Cells that met unique molecular index (UMI) and gene thresholds were included in subsequent analyses. Clustering was performed based on a principal component analysis and visualized using UMAP (McInnes et al., 2018). Differential expression between the clusters was calculated using the negative binomial test (Figures 1 and 2) and Moran’s I tests (Figure 2).

### Bulk RNA-Seq Analysis

Trimming of known adapters and low-quality regions of reads was performed using Fastq-mcf (http://code.google.com/p/ea-utils). Sequence quality control was assessed using FastQC (http://www.bioinformatics.babraham.ac.uk/projects/fastqc/) and RSeQC (Wang et al., 2012). Alignment of the provided samples to the mm9 reference genome was performed using Bowtie 2.2.4 (Langmead and Salzberg, 2012). Reads were assigned to genes using “featureCounts”(Langmead and Salzberg, 2012; Liao et al., 2014), part of the Subread suite (http://subread.sourceforge.net/). Gene-level counts were arrived at using Ensembl gene annotation, in GTF format. We calculated differential expression p-values using edgeR, an R package available through Bioconductor (Robinson et al., 2010). We first filtered out genes where there were not at least two samples with at least 5 (raw) reads. Once these genes are removed, we recalculate the counts per million for each gene (CPM) and filter out any genes with a CPM above 20,000. After excluding these genes, we re-normalize remaining genes using calcNormFactors (TMM) (“weighted trimmed mean of M-values”) in edgeR (Robinson and Oshlack, 2010; Robinson et al., 2010). Calculation of p-values is performed in edgeR for the differential expression between samples. EdgeR uses a negative binomial distribution as a model for expected gene expression. Finally we use the built-in R function “p.adjust” to calculate the FDR (false discovery rate) for each P-value, using the Benjamini-Hochberg method (Benjamini and Hochberg, 1995). We used GO-Elite to generate biological ontology terms (Zambon et al., 2012).

### ATAC-Seq Analysis

Reads were aligned to the mm9 genomic assembly using bowtie2 with options: -X 600 --no-mixed --no-discordant. Duplicate reads were removed using Picard MarkDuplicates (http://broadinstitute.github.io/picard). Peaks were called using macs2 callpeak with options: -p 0.1 --nomodel --shift 100 --extsize 200 -B --SPMR --call-summits. Peaks concordant between at least two of three replicates were considered for further analysis. The clustering is performed using the bioconductor package HOPACH and visualized using pheatmap in R (Laan et al., 2003). The regions are determined by first estimating counts in each of the replicate samples for each of the time-points across a merged set of 307,204 peaks (called by MACS2), then normalizing the counts for differences in sequencing depths and estimating the association with time using the likelihood ratio tests based on negative binomial generalized log-linear model in bioconductor package edgeR. The 10,000 regions represent those with most significant association from the 100,691 regions passed an FDR threshold of 0.05 based on this test. We associated biological processes to these regions using GREAT (McLean et al., 2010). Motif enrichment analysis was completed using HOMER (Heinz et al., 2010).

### ChIP-Seq Analysis

Trimming of known adapters and low-quality regions of reads was performed using Fastq-mcf (http://code.google.com/p/ea-utils). Sequence quality control was assessed using FastQC (http://www.bioinformatics.babraham.ac.uk/projects/fastqc/) and RSeQC (Wang et al., 2012). Alignment of the provided samples to the mm9 reference genome was performed using Bowtie 2.2.4 (Langmead and Salzberg, 2012). Peaks were called using GEM (Guo et al., 2012; Langmead and Salzberg, 2012). Raw read counts per peak were generated with featureCounts (Liao et al., 2014), DEseq2 to normalize read counts, then ChIP signal was normalized to input before using hopach() to cluster peaks with the following settings: clusters=“best”, initord=“clust” (Laan et al., 2003). Region intersects were found using BEDTools (Quinlan and Hall, 2010). Motif enrichment analysis was completed using HOMER (Heinz et al., 2010). Known motifs and sequence logos were generated from *de novo motifs* matched to the JASPAR CORE non-redundant vertebrate database (Heinz et al., 2010; Mathelier et al., 2015) using Tomtom from the MEME suite (Bailey et al., 2009). Significant differences in peak distances from gene groups was determined by Wilcoxon-Mann-Whitney rank sum testCounts were normalized for differences in sequencing depth between samples using upper quartile normalization separately for the ChIP and the input samples of each transcription factor. The normalized counts for the three transcription factor were then combined into one matrix, the top 500 most variable regions based on these normalized counts were determined, and corresponding counts for these regions are used to perform principal component analysis.

### Mathematical Modeling of Transcription Factors Associated with Changing Rate of Gene Expression over Time: Gene Regulation Model for Integrating Changes in Chromatin State with Gene Expression

The net importance of a given transcription factor motif captures the mean fold-change (day 2 versus fibroblast stage) across all genes that gain occurrences of this motif resulting from chromatin changes in a 2kb-500kb neighborhood around their transcription start sites, during this transition versus the mean fold-change across all genes that lose occurrences of this motif during the same transition, after accounting for the effects for all other transcription factor motifs. These were estimated using a targeted maximum likelihood estimation (tmle) as implemented in the tmle package in R that used random forests and generalized linear models as the underlying machine learning algorithms. The targets of a given transcription factor that gain occurrences of the motif are empirically determined as the set of genes whose change in motif association due to the time transition is greater than the 90th quantile of the changes in motif association across all genes. Similarly the targets of a transcription factor motif that loses association are the set of genes whose change in motif association due to the time transition is less than the 10th quantile of the changes in motif association across all genes. (See Supplemental Methods for additional information.)

## SUPPLEMENTAL INFORMATION

Supplemental information includes 6 figures, 10 tables, and an extended methods section describing modeling approach applied in Figure 6.

### SUPPLEMENTAL FIGURES

**Figure S1.**
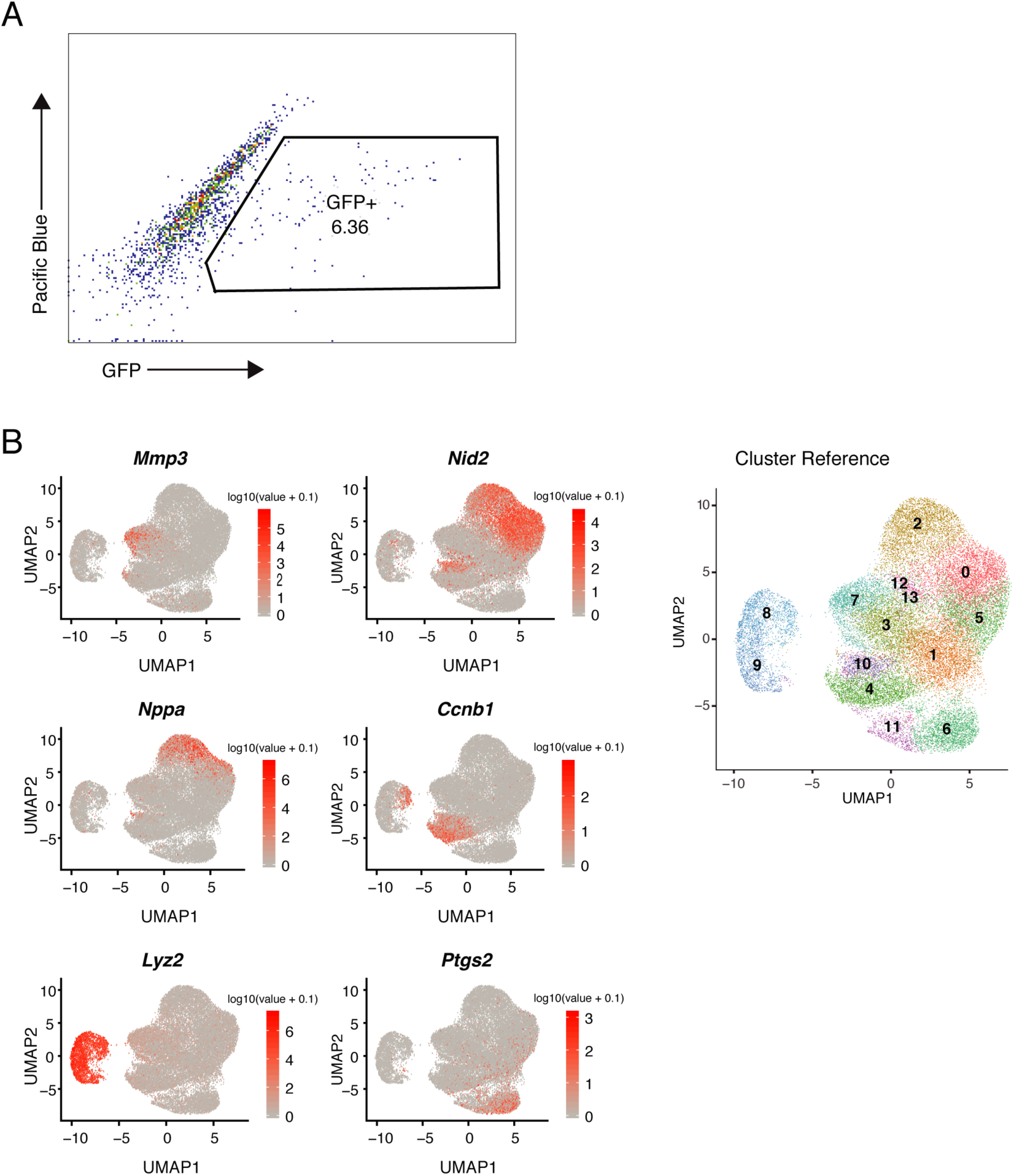
(A) Representative fluorescence-activated cell sorting (FACS) plots for αMHC-GFP+ cells at week 2 of reprogramming. (B) Expression (log10(UMI+0.1)) of marker genes for each cluster visualized in UMAP plots. Numbered clusters from Figure 1B listed at right for reference.

**Figure S2.**
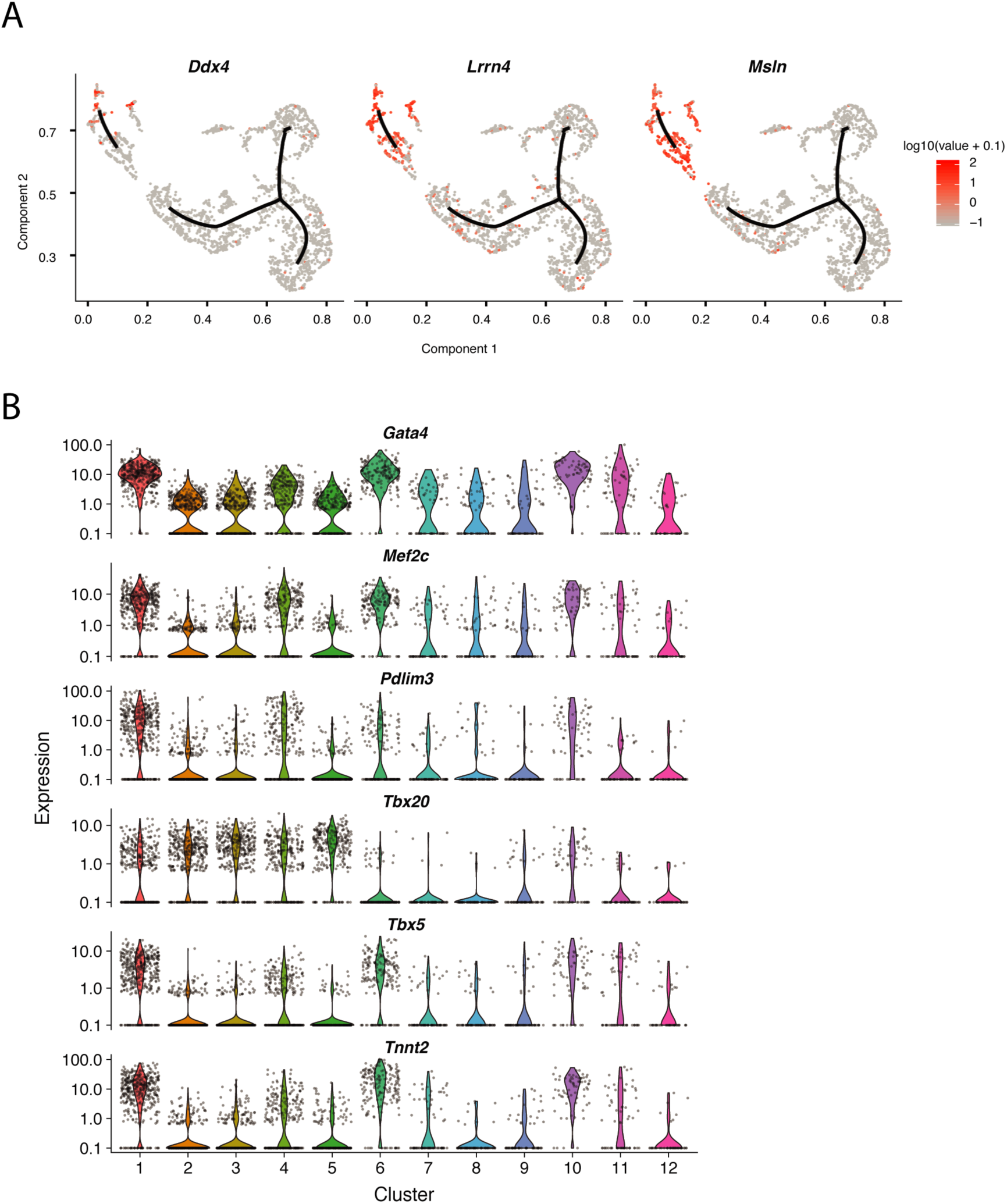
(A) Expression (log10(UMI+0.1)) of epicardial markers visualized in a UMAP trajectory plot. (B) Violin plots representing normalized UMI levels of genes presented in **Figures 2F-G** for all clusters.

**Figure S3.**
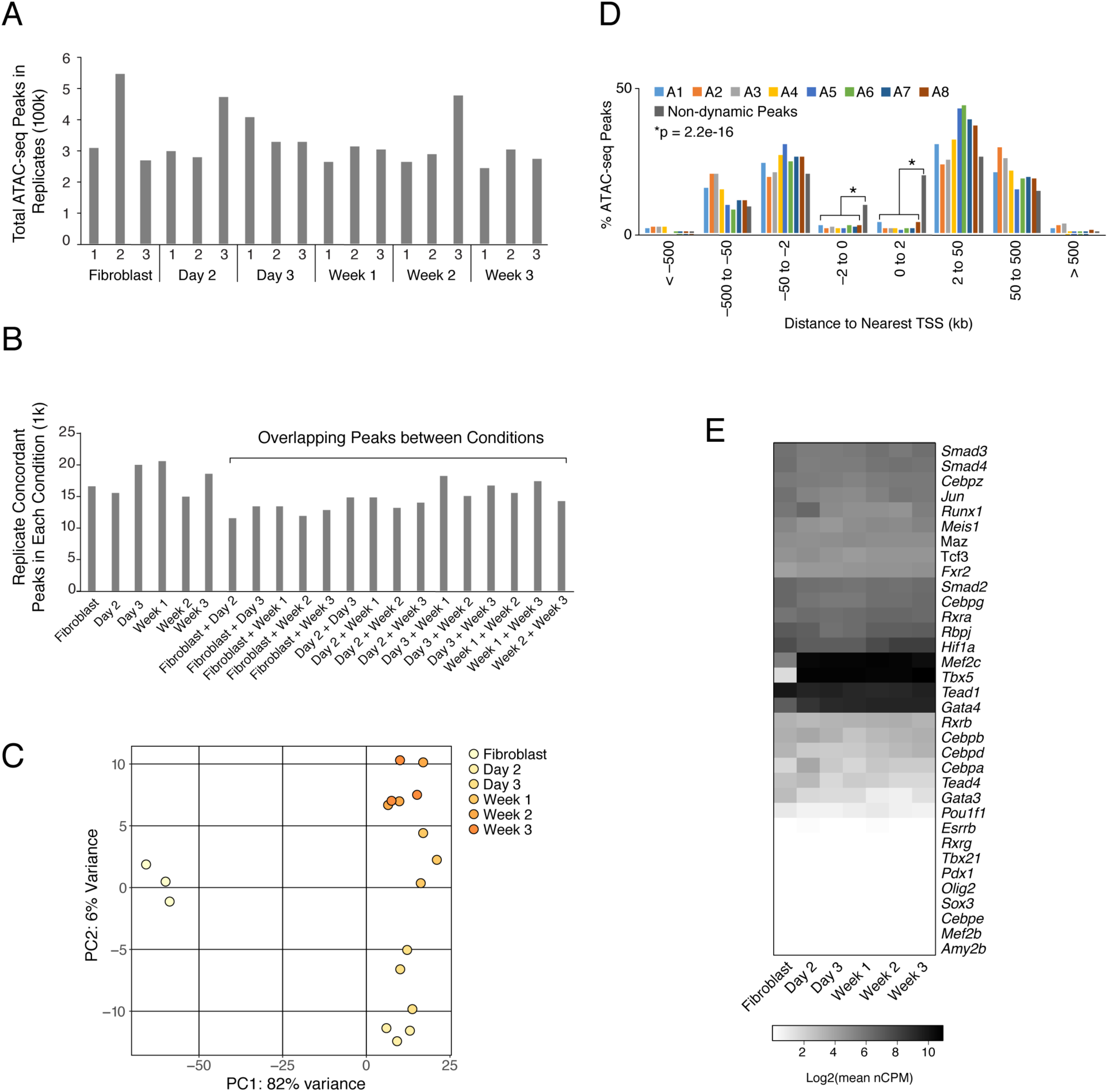
(A) Bar graph indicates total number of ATAC-seq peaks identified for each replicate (1-3). Sorted αMHC-GFP+ cells were assayed at each time point and in sorted dsRed-positive fibroblasts infected with a control dsRed retrovirus. (B) Bar graph indicates total number of concordant ATAC-seq peaks for each condition and total number of replicate concordant peaks overlapping between conditions. Concordant peaks met an FDR threshold of < 0.05 in at least two out of three replicates. (C) Principal component analysis of all ATAC-seq replicates (n=3) for each condition based on 10,000 most differentially accessible regions among all reprogramming time points compared with fibroblasts. (D) Histogram shows distribution of regions within each dynamic cluster (A1-A8) and in a set of non-dynamic regions that are stably accessible in fibroblasts and in all reprogramming conditions, binned based on the distance from each peak to the nearest transcription start site (TSS). Dynamic regions were underrepresented within 2kb of a TSS compared with non-dynamic regions accessible in all conditions (Fisher’s p-value = 2.2e-16 for both up- and downstream bins). (E) Heatmap displays expression levels for transcription factors with motifs with highest enrichment in dynamic accessibility clusters (A1-A8) compared with non-dynamic region set stably accessible in all conditions. Values indicate log2 of the mean normalized CPM across all replicates (n=3) from bulk RNA-seq completed on sorted αMHC-GFP+ cells collected at each time point during reprogramming.

**Figure S4.**
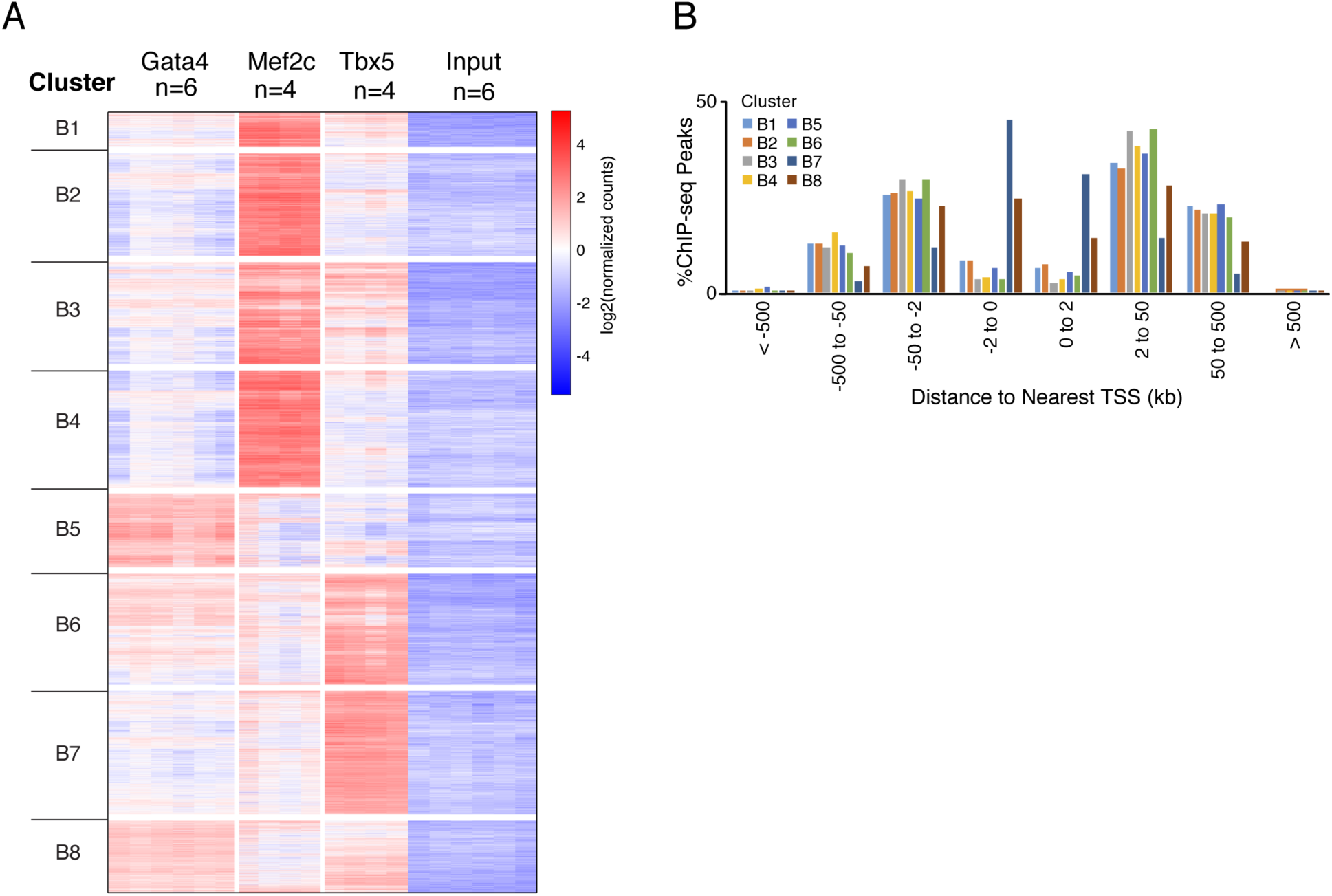
(A) Heatmap displays tag densities of ChIP-seq sample and inputs, for all replicates. Counts were normalized for differences in sequencing depth between samples using upper quartile normalization separately for the ChIP and the input samples of each transcription factor. Columns represent independent replicates. Rows retain order shown in Figure 4B. (B) Bar plot displays binned distribution of ChIP-seq peaks from nearest transcription start site (TSS). We identified a lower proportion of dynamic regions (all clusters) within 2kb up- and downstream of the nearest TSS compared with stably accessible, non-dynamic regions (Fisher odds ratio = 0.2 and 0.07, respectively; p-value = 2.2e-16 for both comparisons).

**Figure S5.**
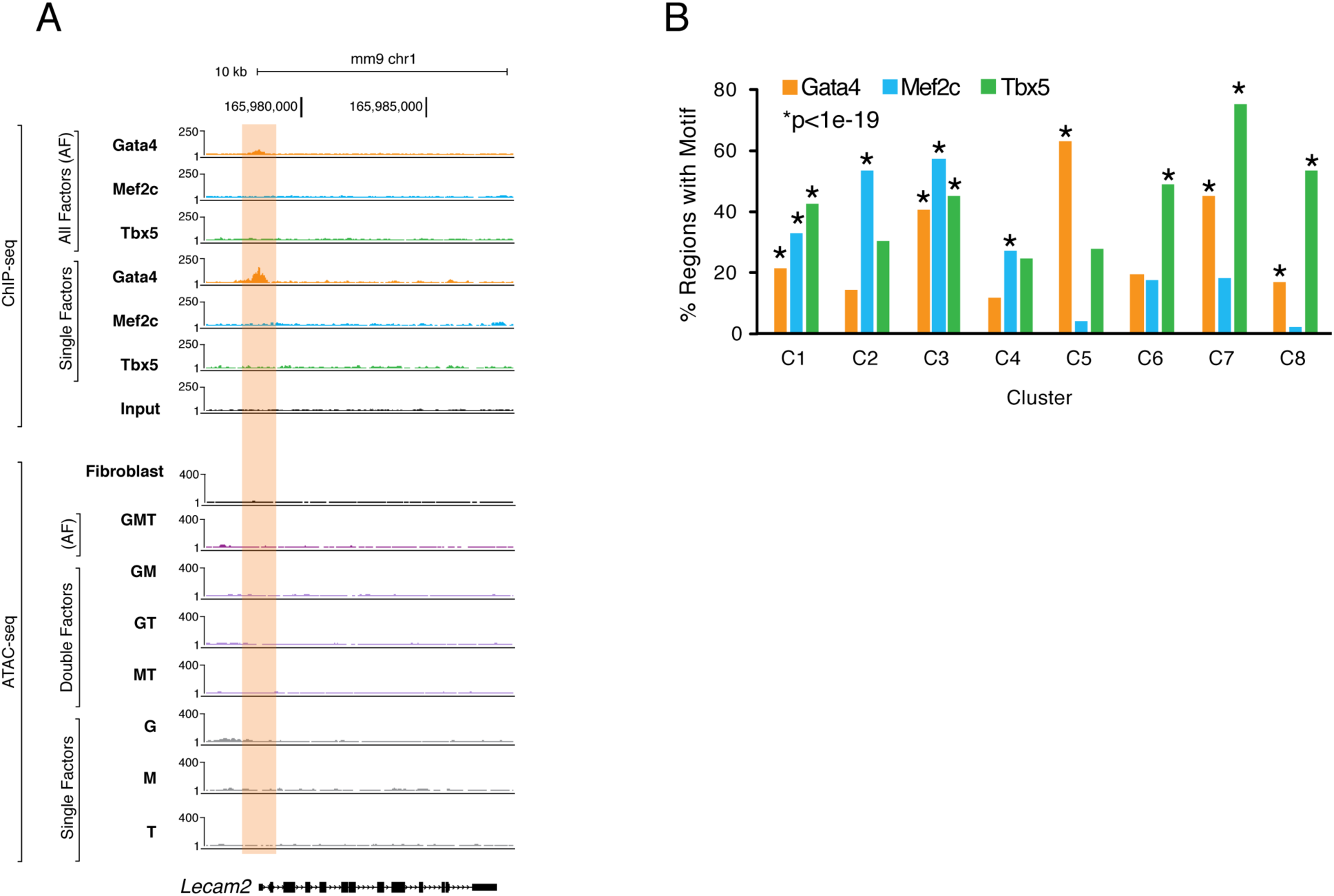
(A) Browser tracks display ChIP-seq in single factor and all factor conditions, and ATAC-seq signal in single factor, double factor, and all factor conditions, near leukocyte-endothelial cell adhesion molecule 2 (*Lecam2*). Track height was normalized to sample read depth within each assay. Orange rectangle highlights region from cluster 5 with Gata4 binding in the single factor condition which is ameliorated when all factors are present. (B) Bar graph indicates percentage of regions within each cluster (C1-C8) that contain Gata4 (orange), Mef2c (blue), or Tbx5 (green) motifs. P-values were calculated using the cumulative hypergeometric distributions of motif occurrence in clustered region set compared with motif occurrence in non-dynamic regions stably accessible in fibroblasts and reprogramming conditions. Full motif enrichment results available in **Table S8**.

**Figure S6.**
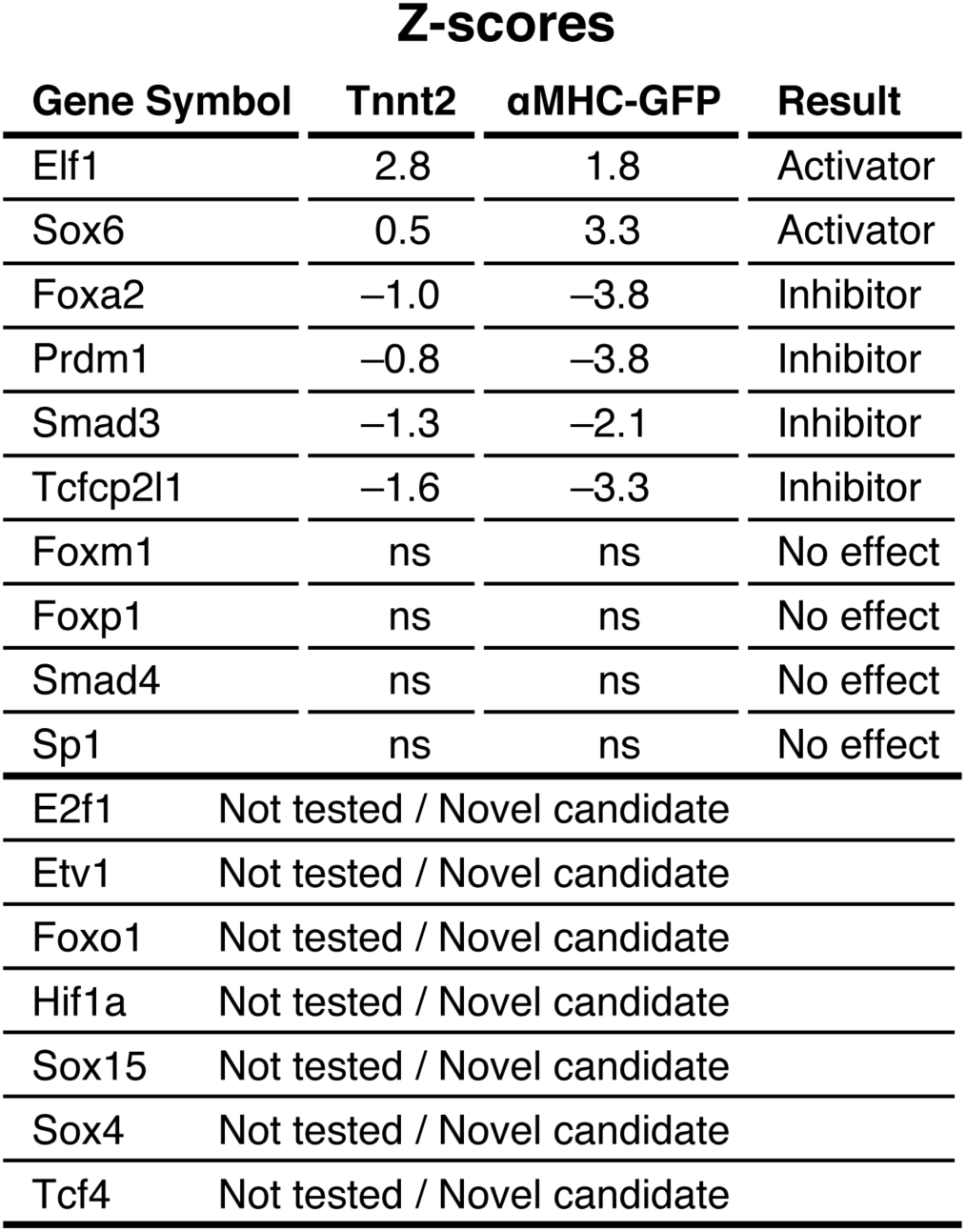
Reprogramming Outcomes Following Candidate Transcription Factor Overexpression. Table lists selected z-score results of reprogramming screen completed by (Zhou et al., 2017) for overexpression of candidate transcription factor predicted in this study to enhance or serve as barriers of cardiac reprogramming in a cooperative manner, highlighted in **Figure 6D**.

### SUPPLEMENTAL TABLES

**Table S1. Related to Figure 1. Differential Gene Expression for Each Cluster.** Contains average log fold changes and p-values for all differentially expressed genes from negative binomial tests used to determine identities of populations in clusters in **Figure 1B**, and designates top 20 marker genes per cluster used to generate **Figure 1E**.

**Table S2. Related to Figure 2. Differential Gene Expression for Each Cluster.** Contains average log fold changes and p-values for all differentially expressed genes from negative binomial tests used to determine identities of populations in clusters in **Figure 2A**, and results of Moran’s I tests, including top two marker genes per cluster used in **Figure 2C**. Also contains average fold changes and p-values for all differentially expressed genes from pairwise tests of cluster 2 vs cluster 1, cluster 1 vs cluster 4, and cluster 4 vs cluster 8.

**Table S3. Related to Figure 3.** Table containing 100,691 regions dynamic between fibroblasts and αMHC-GFP+ cells collected at reprogramming time points day 2, day 3, week 1, week 2, and week 3.

Table S4. Related to **Figure 3**. Known motif enrichment results comparing regions in each cluster to stably accessible background region set.

**Table S5. Bulk RNA-seq Gene Expression during Reprogramming.** Table includes raw read counts and normalized CPM values for each replicate for all genes expressed at an average value nCPM > 1 in fibroblasts or αMHC-GFP+ cells over time course. Includes differential expression results between conditions.

Table S6. Related to **Figure 3**. Gene Ontology Results from Regions Clustered by Chromatin Accessibility Changes over Time Using GREAT Analysis.

Table S7. Related to **Figure 4**. Known Motifs Enriched in Regions Clustered by Gata4, Mef2c, Tbx5 Occupancy and Chromatin Accessibility at Day 2 of Reprogramming.

Table S8. Related to **Figure 5**. Known Motifs Enriched in Regions Clustered by Gata4, Mef2c, Tbx5 Occupancy and Chromatin Accessibility during Combinatorial Exogenous Expression of Individual Factors.

Table S9. Related to **Figure 5**. Fold Change in ChIP-seq Signal over Input and ATAC-seq Signal over Fibroblast Control for All Regions in **Figure 5C**.

**Table S10. Related to Figure 6. Total Predicted Co-regulatory Interactions of Transcription Factor Candidates.** Table includes total number of cofactors predicted by model for candidate regulators of reprogramming, with totals listed separately for predicted interactors with positive and negative net importance scores.

## SUPPLEMENTAL METHODS

### Proteins/transcription factors associated with changing rate of gene expression over time

#### Gene regulation model for integrating changes in chromatin state with gene expression

A model is constructed for the differences in the rates of change of mRNA levels for each gene between two consecutive time-points as a function of changes in sequence composition of differentially open and closed regions between these time-points in a *cis*-region for the gene. Only changes in sequence composition associated with known transcription factors are used in this analysis.

Let *M* denote the number of genes (this corresponds to the set of genes whose mean expression is associated with time or the day of reprogramming). Let *T_t_* denote the *t^th^*time-point, *t* ∈ {0,1, 2, 3, 4, 5}, where gene expression and chromatin state is assayed (*T_0_ =* 0 [Fibroblast stage before the GMT vectors are added], *T_1_ =*2 [day 2], *T_2_ =*3 [day 3], *T_3_ =*7 [week 1], *T_4_ =*14 [week 2] and *T_5_ =*21 [week 3]).

Let *X_t,i_* denote the *log2* mean (across the 3 replicates) normalized (across all replicates over time) expression of gene *i* at time *t* . Let *Y_t,i_* denote the rate of change of the logarithm of the mean of expression of gene *i* at time *t*.

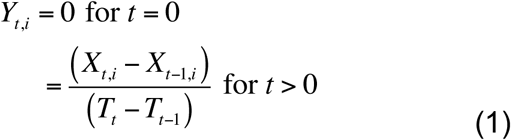

Let Δ*Y_t_*_,*i*_ denote the difference between the rates of change of the *log2* expression of gene *i* at time *t* and at time *t-1*.

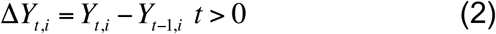

Assume that the changes in rate of change of the expression of gene *i* between time *t* and time *t-1* can be explained by some subset of *N* sequence motifs. Each sequence motif is associated with a protein/transcription factor. Let 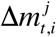 denote the change in the strength of association of motif *j*, *j* ∈ {1, 2,!, *N*}with gene *i* between time *t* and time *t-1* resulting from the differential opening and closing of chromatin.

One model of gene expression regulation across all genes between times *t-1* and *t* (denoted by *F_t_*) is assumed.

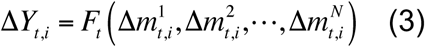

The functional form of *F_t_* is non-parametrically identified using a supervised learning approach (see next sections).

#### Biophysical motivation for the modeling approach

The above gene regulation model has a biophysical motivation. The rate of change of the log of the expression of a given gene at a given time is the difference between the rate at which it is transcribed and the rate at which the corresponding mRNA decays. The rate of transcription of this gene is a (unknown) function of the strengths of association of transcription factors/proteins with this gene at this time-point. The rate of decay of the log of the expression of the gene is a fixed constant (independent of time) if one assumes a first-order rate of decay for the corresponding mRNA. Therefore the differences in the rate of change of the log of the expression of gene at the two time-points should be a function of the difference in the strengths of association of the transcription factors between these time-points. Note the analysis assumes the same model, *F_t_* (Equation 3), for all genes. This is a simplification necessitated by grossly smaller number of samples (order 1) versus the number of possible interacting motifs/transcription factors (order 100) and will result in the identification of modes of regulation that is apparent across a relatively large proportion of genes. On the other hand this simplification has the advantage of not directly requiring the concentrations of the regulating transcription factors corresponding to over-represented sequence motifs.

#### Identification of the list of proteins/transcription factors

The list of sequence motifs/proteins/transcription factor to use for the model between consecutive time-points are identified using the open chromatin regions for each replicate ATAC-seq sample at these time-points (see MACS2 section for peak calling). The open chromatin regions at each time-point by combining the regions across replicates (using *bedops –everything* [1]). The *findMotifsGenome.pl* function (using the options *–size given* and hypergeometric enrichment scoring) in *Homer* [2] is used to identify motifs enriched (p-value < 1e-1000) at time *t*, using the open chromatin regions at time t as foreground and the open chromatin regions at time *t-1* as background. Similarly, this function is used to identify motifs enriched at time *t-1* using the open chromatin regions at time, *t* as background. The list of proteins used in the analysis at time *t* is the union of the motifs from the two above enrichment analyses.

#### Association strength of a motif with a gene

The location of each of the above identified motifs in the open chromatin regions at each time-point is obtained using the *–find* option of the *findMotifsGenome.pl* function. Assume that there are 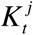 motif locations of motif *j* in the open chromatin regions at time Then the strength of association of motif *j* with gene *i* is given by,

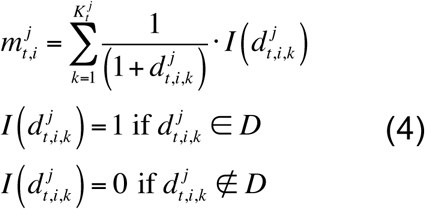

where 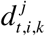 is the distance of the k^th^ location of the motif from the transcription start site (TSS) of gene *i. D* corresponds to a distance domain. In the analysis, two distance domains are considered – (0, 2kb) and (2kb, 500kb) corresponding to promoter and potential distal enhancer associations.

The change in motif gene association is then defined as,

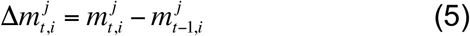

#### Validation of the gene regulation model

The gene regulation model stated in Equation (3) is fit across a set of genes using the random Forests [3] based supervised learning approach. This is done using the *rfsrc* function that is part of the *randomForestSRC* package [4] in R[5]. The set of genes whose mean expression is associated with the time or the day of reprogramming is randomly divided into ten groups. The data for the genes corresponding to nine of the ten groups are used to learn the model given in Equation (3). The correlation between the observed Δ*Y_t_*_,*i*_ for the set of the genes in the remaining group and the predicted Δ*Y_t_*_,*i*_ for these genes using the model learnt is computed.

#### Importance of transcription factors/proteins in explaining changes in gene expression

The importance of each of the transcription factors in explaining changes in rate of change of expression across all the genes between time-point *t-1* and *t*, is defined here. This is followed by its estimation procedure.

In words, the importance of a given transcription factor at a given time-point, *t* is defined as the change in the mean difference in the rates of changes of expression of genes which is associated with this transcription factor from the mean difference in the rates of changes of expression of genes which are not associated with this transcription factor after accounting for effects from all other transcription factors on this difference.

Denote the set of positive values of 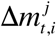 by,

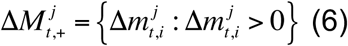

the second of negative values of 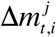 by

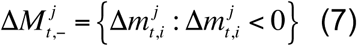

and the set of absolute values of 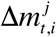, by

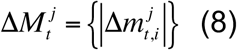

Define 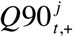 as the 90^th^ quantile of 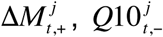 as the 10^th^ quantile of 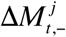 and 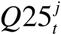 as the 25^th^ quantile of 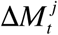.

Denote 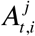 as a binary variable that is equal 1 if motif *j* is associated with gene *i* at time. *t*.

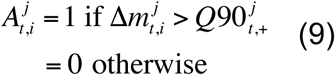

Denote 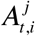 as a binary variable that is equal -1 if motif *j* is associated with gene *i* at time *t*-1

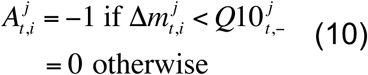

Denote 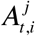 as a binary variable that is equal 2 if motif *j* is not associated with gene *i* at either time-point.

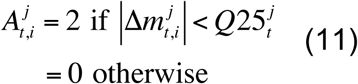

Let 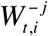 denote the vector of changes in motif association with gene *i* across all motifs except motif *j*.

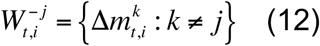

Then the marginal mean difference in rate of change of expression between time-points *t* and *t-1* across genes associated with motif j at time *t* is defined as,

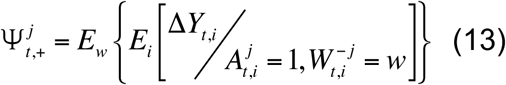

The symbol *E* denotes expectation while its subscript denotes the values over which the expectation is taken. The marginal mean difference in rate of change of expression between time-points *t* and *t-1* across genes associated with motif j at time *t-1* is defined as,

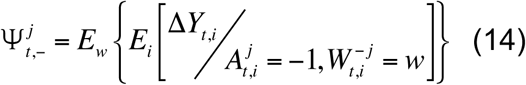

The marginal mean difference in rate of change of expression between time-points *t* and *t-1* across genes not associated with motif j at either time is defined as,

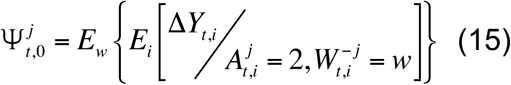

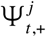, 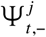 and 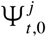 are estimated using the targeted Maximum Likelihood Estimation (tMLE) approach [6]using the *tmle* package [7]in R. Random forests and Generalized Linear Models (*glm*) are the two model specified for use by the SuperLearner [8] for estimation in the *tmle* function.

The importance of motif j at time point t is then defined as,

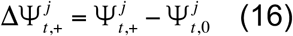

The importance of motif j at time point t-1 is defined as,

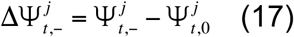

The statistical significance of 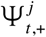 and 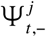 are determined from the standard errors of 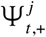, 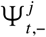 and 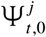 estimated using the *tmle* function. The significant motifs for time-points *t* and *t-1* are identified using a Bonferroni-defined threshold of 0.05/(2N), where *N* is the number of identified enriched motifs in the open chromatin regions at these time-points.

The net importance of each motif is then defined as,

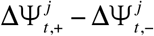

The bar plots in Figure 6b represent the net importance values of the transcription factors. The net importance of a given transcription factor motif captures the mean fold-change (day 2 versus fibroblast stage) across all genes that gain occurrences of this motif resulting from chromatin changes in a 2kb-500kb neighborhood around their transcription start sites, during this transition versus the mean fold-change across all genes that loose occurrences of this motif during the same transition, after accounting for the effects for all other transcription factor motifs. While both positive and negative scores indicate associations with dynamically expressed genes, a positive score for a transcription factor would be consistent with an activating influence on gene expression while a negative score with a repressing influence on gene expression.

